# Concurrent depletion of Vps37 proteins evokes ESCRT-I destabilization and profound cellular stress responses

**DOI:** 10.1101/2020.07.02.183954

**Authors:** Krzysztof Kolmus, Purevsuren Erdenebat, Blair Stewig, Ewelina Szymańska, Krzysztof Goryca, Edyta Derezińska-Wołek, Anna Szumera-Ciećkiewicz, Marta Brewińska-Olchowik, Katarzyna Piwocka, Monika Prochorec-Sobieszek, Michał Mikula, Marta Miączyńska

## Abstract

Molecular details of how endocytosis contributes to oncogenesis remain elusive. Our *in silico* analysis of colorectal cancer (CRC) patients revealed stage-dependent alterations in the expression of 113 endocytosis-related genes. Among them transcription of the Endosomal Sorting Complex Required for Transport (ESCRT)-I component *VPS37B* was decreased in the advanced stages of CRC. Expression of other ESCRT-I core subunits remained unchanged in the investigated dataset. We analyzed an independent cohort of CRC patients showing also reduced *VPS37A* mRNA and protein abundance. Transcriptomic profiling of CRC cells revealed non-redundant functions of Vps37 proteins. Knockdown of *VPS37A* and *VPS37B* triggered p21-mediated inhibition of cell proliferation and sterile inflammatory response driven by the Nuclear Factor (NF)-κB transcription factor and associated with mitogen-activated protein kinase signaling. Co-silencing of *VPS37C* further potentiated activation of these independently induced processes. The type and magnitude of transcriptional alterations correlated with the differential ESCRT-I stability upon individual and concurrent Vps37 depletion. Our study provides novel insights into cancer cell biology by describing cellular stress responses that are associated with ESCRT-I destabilization, which might occur in CRC patients.

**SUMMARY STATEMENT:** Endosomal Sorting Complex Required for Transport (ESCRT)-I destabilization upon concurrent depletion of Vps37 proteins is linked to the activation of sterile inflammatory response and cell growth inhibition.

## INTRODUCTION

Genetic alterations induce the reprogramming of intracellular signaling, which is a driving force of tumorigenesis. The duration of signal transduction is dependent on endocytosis (Floyd and De Camilli, 1998; Mosesson et al., 2008; Schmid, 2017). Some mechanisms for tumor cell-specific changes in the activity of endocytic machinery components that affect intracellular signaling have been already identified (Barbieri et al., 2016; Di Fiore and von Zastrow, 2014; Mellman and Yarden, 2013). For instance, many tumors exhibit deregulated expression of the ubiquitination machinery and small GTPases that control the rate of receptor degradation and recycling, respectively (Porther and Barbieri, 2015). However, despite the abundance of publicly available data, such as those deposited in The Cancer Genome Atlas [TCGA] (Weinstein et al., 2013), little has been done to systematically analyze the expression of receptor trafficking regulators in tumors and across tumor stages. This knowledge could potentially facilitate patients’ stratification for treatment with bioengineered macromolecules delivered through receptor-mediated endocytosis (Tashima, 2018).

An important group of trafficking regulators constitute four sequentially acting Endosomal Sorting Complexes Required for Transport (ESCRT-0, ESCRT-I, ESCRT-II, and ESCRT-III) and accessory proteins, among others Vps4A and Vps4B. The ESCRT machinery mediates receptor degradation not only by recognition and local clustering of ubiquitinated cargo on endosomes but also through membrane deformation and scission to form intraluminal vesicles (ILVs). Many rounds of ILV formation create multivesicular bodies that fuse with lysosomes leading to cargo degradation. In addition, ESCRTs contribute to cytokinesis, autophagy, virus budding, exovesicle release, and repair of plasma and intracellular membranes (Hurley, 2015; Olmos and Carlton, 2016; Szymanska et al., 2018; Vietri et al., 2020). Despite the well-established roles of ESCRT components in maintaining cell homeostasis, much less is known about their contribution to tumorigenesis (Alfred and Vaccari, 2016; Gingras et al., 2017; Mattissek and Teis, 2014) and the underlying molecular mechanism has been clarified only in a couple of cases (Manteghi et al., 2016; Sadler et al., 2018). For instance, we demonstrated that the expression of *VPS4B*, encoding ESCRT-associated ATPase, is decreased in colorectal cancer (CRC) and *VPS4B*-deficient cells are critically dependent on the Vps4A protein. This synthetic lethality between *VPS4* paralogs triggers stress-associated sterile inflammatory response and immunogenic cell death and thus may be used as a basis for personalized therapy (Szymanska et al., 2020).

ESCRT-I is a heterotetramer composed of three core components (Tsg101, Vps28 and one of four Vps37 family members) and a single auxiliary protein (UBAP-1, Mvb12A or Mvb12B) (Stefani et al., 2011; Wunderley et al., 2014). At least under some conditions two of its subunits (Tsg101 and Vps37A) have been identified as putative tumor suppressors (Li and Cohen, 1996; Moberg et al., 2005; Xu et al., 2003). In parallel, high-throughput screens for cancer vulnerability within the DepMap project (Behan et al., 2019) demonstrated that multiple cancer cell lines display a reduced fitness upon *TSG101* knockout, whilst the effect of perturbed expression of *VPS28, VPS37A, VPS37C* or *UBAP1* genes is cell type-dependent.

Tsg101 and Vps37A are not only regulators of vesicular trafficking but also other biological processes, such as transcription and autophagy (Bache et al., 2004; Bishop et al., 2002). Transcriptomic analysis of Tsg101-depleted cancer cells revealed increased expression of the prototypical Nuclear Factor-ĸB (NF-ĸB)-dependent genes without exogenous stimulation (Brankatschk et al., 2012). We dissected the molecular basis of this phenomenon showing that the absence of Tsg101 or Vps28 led to the accumulation of ligand-free cytokine receptors on endosomes because of disturbed sorting into ILVs and degradation of cargo. The proximity of accumulated receptors on endosomes evoked their oligomerization to trigger NF-ĸB signaling (Banach-Orlowska et al., 2018; Maminska et al., 2016). However, none of the Vps37 proteins incorporated into ESCRT-I was identified as a genuine regulator of the NF-ĸB pathway.

NF-ĸB is a family of ubiquitously expressed transcription factors, whose activation is a hallmark of inflammation often associated with cancer (Taniguchi and Karin, 2018). These transcription factors mediate also other biological processes, such as proliferation (Zhang et al., 2017). There are two interconnected NF-ĸB signaling cascades. The canonical NF-ĸB pathway culminates in the phosphorylation and degradation of the IκBα inhibitor, that allows an active p65-p50 NF-ĸB dimer to translocate into the nucleus. The non-canonical signaling cascade marks the p100 NF-ĸB precursor for proteasomal processing to p52 to form transcriptionally active complexes with RelB (Hayden and Ghosh, 2008). Although endocytic trafficking and NF-ĸB inflammatory signaling are important in carcinogenesis, the molecular links between them are poorly studied. Here, we systematically analyzed the expression of endocytosis regulators across stages of CRC using publically available data and found decreased expression of *VPS37* paralogs. As no genome-wide expression studies have explored the cellular consequences of individual and concurrent depletion of Vps37 family members, we investigated their roles focusing on the processes related to cell growth and inflammatory response. Our findings reveal the importance of *VPS37* paralogs in orchestrating cell homeostasis through maintaining the stability of ESCRT-I.

## RESULTS

### Expression of *VPS37B* is decreased in advanced colorectal cancer

CRC is a leading cause of cancer-associated deaths worldwide as it is often diagnosed in advanced stages when patients display clinical symptoms (Siegel et al., 2018). Aberrant endosomal trafficking in CRC has been linked to adverse phenotype and resistance to therapies (Gargalionis et al., 2015). In order to gain insight into transcriptional changes of genes involved in endosomal trafficking in CRC, we mined the TCGA data against a custom-made list (Table S1) of components whose biological function was related to endocytic transport. Matched normal and cancer transcriptomic samples of human CRC cohorts (colon adenocarcinoma [COAD] and rectal adenocarcinoma [READ]) with available clinicopathological information (31 patients in total) were divided based on tumor staging to early (Stage I and II; 19 patients) or advanced (Stage III and IV; 12 patients) disease stage pools (Fig. 1A). Out of the 445 endocytic genes tested, 410 genes fulfilled normalization criteria of the present analysis (see Materials and Methods). We observed differential gene expression of 113 genes at different stages of CRC when compared to levels transcribed in matched healthy colon tissue (Fig. 1B). Differential expression of 20 genes was unique for the early stages (Table S2). 21 genes were differentially expressed in the advanced stages of tumorigenesis (Table S3). In addition to decreased mRNA abundance of *VPS4B*, which we studied before (Szymanska et al., 2020), we detected reduced expression of an ESCRT-I component – *VPS37B –* in the advanced stages of CRC (Fig. 1C).

**Table 1.**
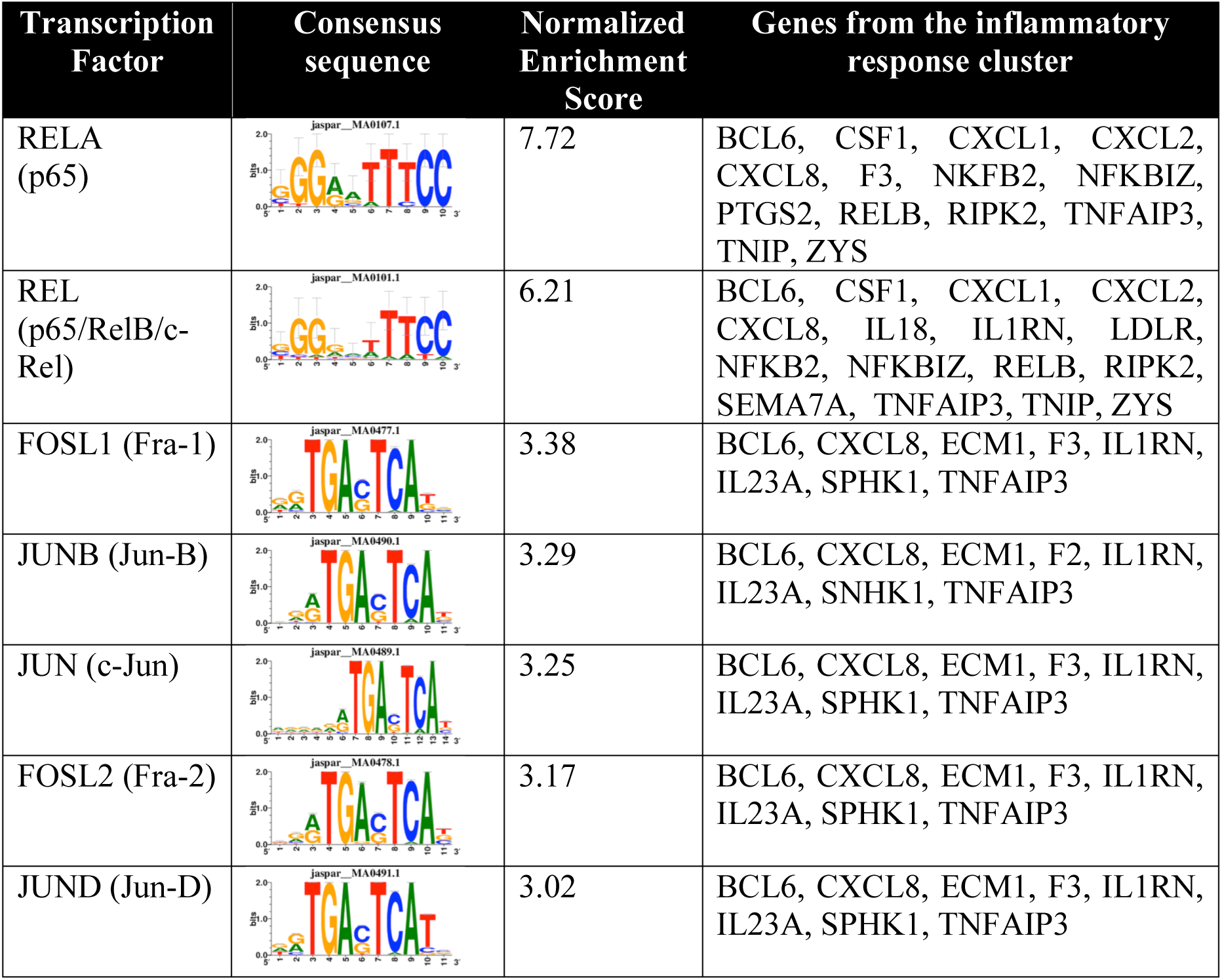
Transcription factors driving expression of genes belonging to the inflammatory response cluster. Transcriptional motif-enrichment analysis of the inflammatory response gene cluster identified in our RNA-Seq analysis was performed using RcisTarget. A region of 500 bp upstream and 100 bp downstream to the transcription starting site was investigated with the TFBS matrices from the JASPAR database.

**Fig. 1.**
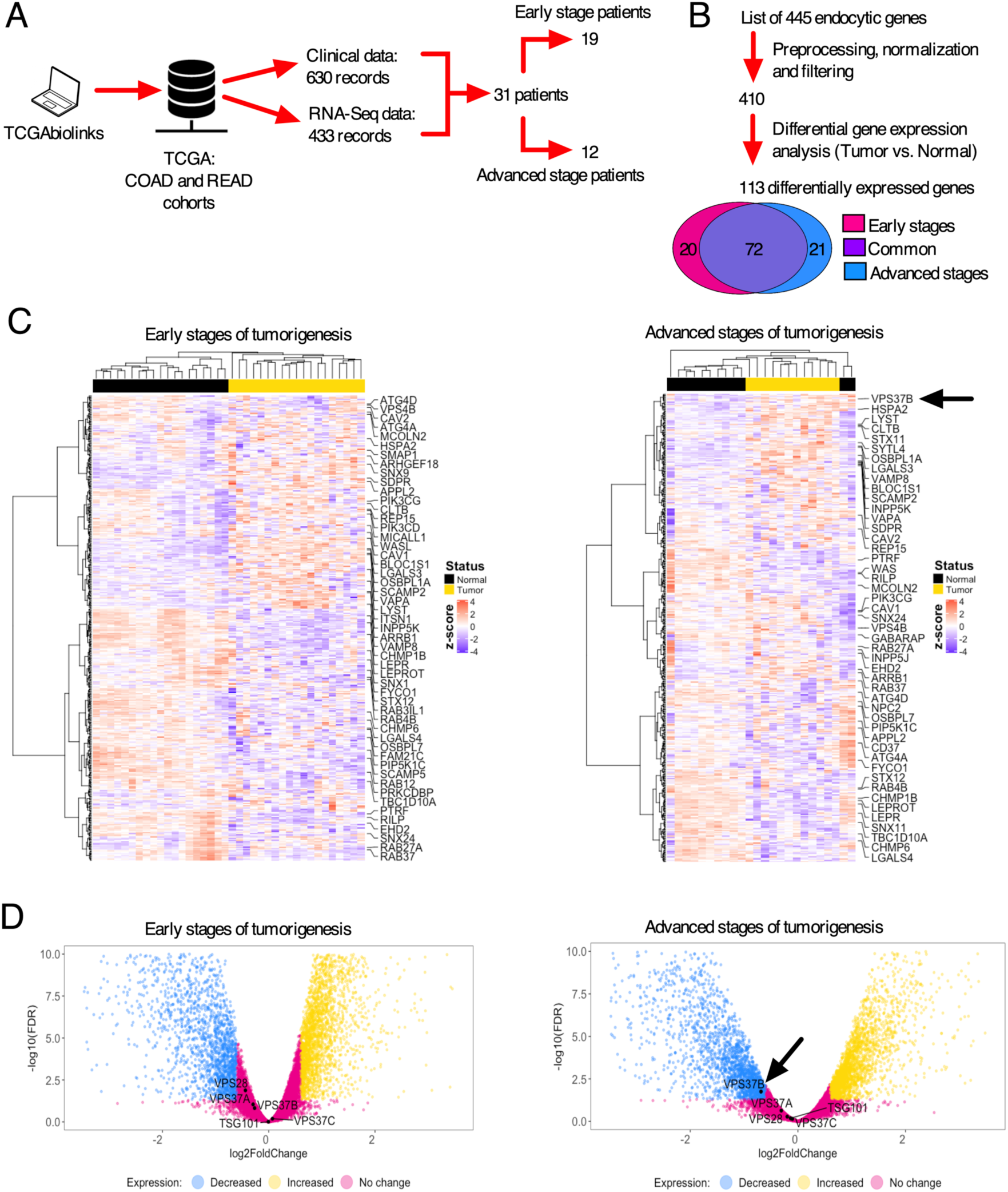
Expression of *VPS37B* is decreased in advanced stages of colorectal cancer. (A) Scheme of data mining process of The Cancer Genome Atlas (TCGA) database with a focus on the COAD and READ cohorts using the TCGAbiolinks package. (B) Scheme showing the number of genes in consecutive stages of analysis and Venn diagram of differentially expressed genes related to endocytic transport in patients at the early (stage I and II) and advanced stages (stage III and IV) of CRC. (C) Heatmaps visualizing the expression of the genes with decreased expression in early stages and advanced stages of CRC. Columns are samples from normal tissue or tumor. Rows are transcripts. (D) Volcano plots visualizing the expression of ESCRT genes in early and advanced stages of CRC. Genes with increased and decreased expression are those with False Discovery Rate (FDR) < 0.05 and log2FoldChange ≥ 0.6 and ≤ −0.6, respectively.

Since we and others had demonstrated that two out of three core ESCRT-I components, namely Tsg101 and Vps28, restrict NF-κB-dependent transcription (Brankatschk et al., 2012; Maminska et al., 2016), we analyzed expression of genes encoding the core ESCRT-I subunits in normal colorectal tissue samples from the TCGA datasets. Using the transcripts per million (TPM) metrics to normalize expression data with respect to gene length and sequencing depth, we observed that colorectal tissue and its cancer counterpart expressed high levels of *TSG101, VPS28, VPS37B*, followed by comparable levels of *VPS37A* and *VPS37C*, and negligible levels of *VPS37D*. Differential expression analysis of CRC samples compared against matching healthy tissue controls revealed that transcription of *VPS37B* tended to be decreased in the early stages of CRC, and was reduced 1.61-fold in the advanced stages (Table S4). Expression of *VPS28* was slightly decreased in the early stages of tumorigenesis (Table S4). On the other hand, levels of *TSG101, VPS37A* and *VPS37C* were stably expressed across tumor stages (Fig. 1D, Table S4).

In summary, expression of *VPS37B* is decreased during progression from early to advanced stages of CRC, whilst transcription of the remaining ESCRT-I components is unchanged.

### mRNA and protein abundance of *VPS37A* and *VPS37B* paralogs is decreased in CRC patient cohorts

Since the expression of *VPS37B* was decreased in samples deposited in the TCGA database and *VPS37A* mRNA and protein abundance was previously shown to be reduced in CRC patients (Chen et al., 2018; Miller et al., 2018; Vasaikar et al., 2019), we performed qRT-PCR analysis of *VPS37* mRNA levels using an independent set of CRC samples from our previous study (Mikula et al., 2011; Skrzypczak et al., 2010). We observed a significant decrease in *VPS37B* and *VPS37A* mRNA abundance in adenocarcinoma (Fig. 2A).

**Fig. 2.**
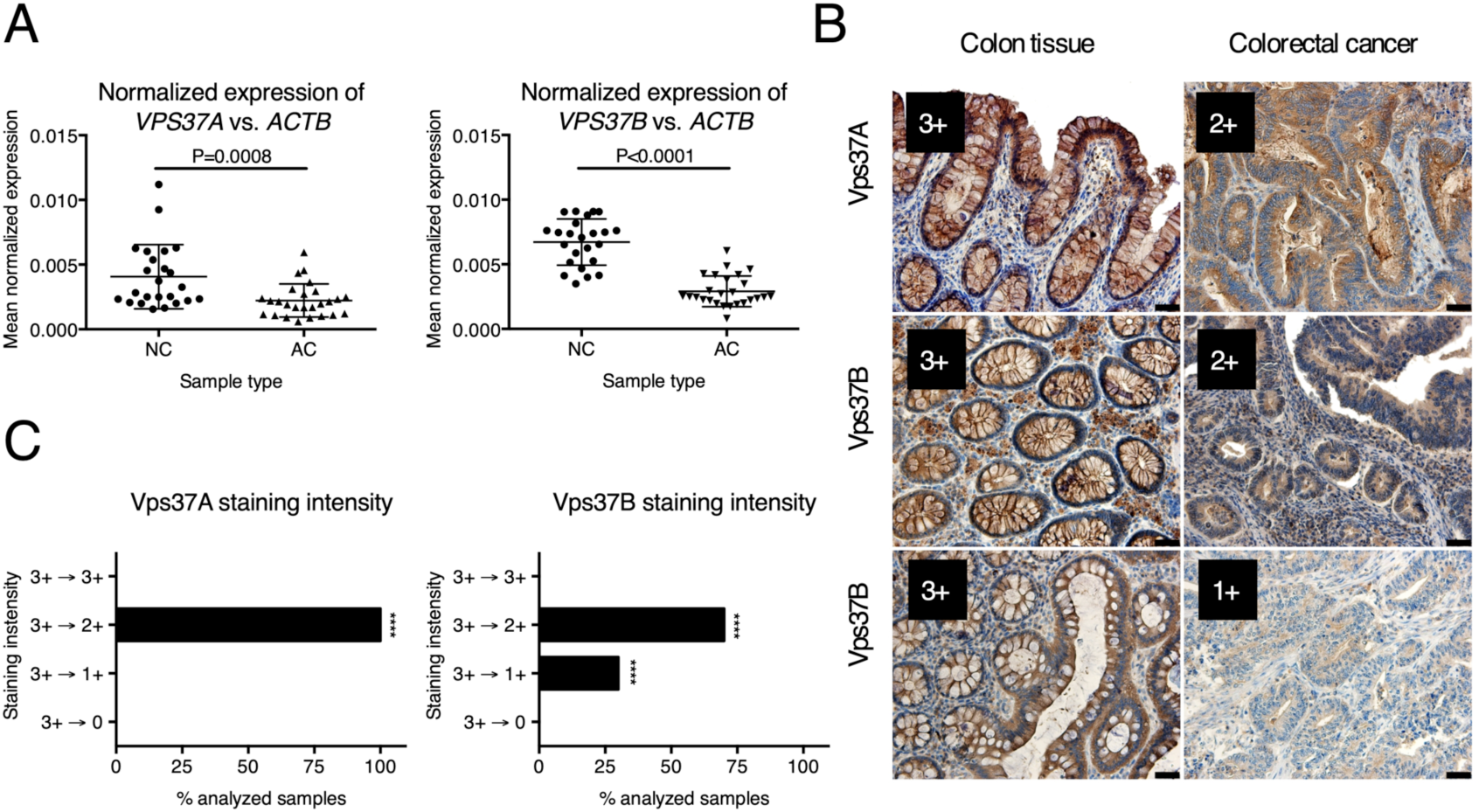
Abundance of *VPS37A* and *VPS37B* paralogs is decreased in a treatment-naïve cohort of CRC patients. (A) qRT-PCR analysis of *VPS37A* and *VPS37B* mRNA levels in normal colon (NC, n=24) and adenocarcinoma (AC, n=26) samples. Middle lines are means and whiskers are standard deviation. Differences were analyzed using the Mann-Whitney U test. (B) Examples of IHC staining of Vps37A and Vps37B in normal colon and matched CRC samples as an illustration of the scoring system used for the evaluation presented in (C). Scale bars: 200 µm. (C) Comparative analysis of Vps37A and Vps37B IHC staining performed in pairs of normal colon and matched CRC samples (n=100): 3+ – very intense staining, 2+ – medium intense staining, 1+ – weak staining, 0 – no staining. Statistical significance was assessed against healthy tissue using Fisher’s exact test \*\*\*\**P*<0.0001.

To assess whether transcriptional alterations at mRNA level correlate with the diminished abundance of Vps37 proteins in CRC, we performed immunohistochemistry (IHC) staining of Vps37A and Vps37B in an array setup consisting of 100 pairs of treatment-naïve primary CRC samples and non-cancerous colon tissue using specific antibodies (Fig. S1A-E). We evaluated the tissue arrays using a semi-quantitative scoring method based on staining intensity.

Both Vps37A and Vps37B displayed strong cytoplasmic staining in normal colon tissue (Fig. 2B). Out of the 100 investigated patient samples, protein staining of Vps37A was decreased to the medium intensity level (3+ --> 2+) in cancerous tissue of all examined patients. Vps37B protein staining was decreased to the medium intensity level (3+ --> 2+) in 70% of patients and to weak intensity levels (3+ --> 1+) in 30% of patients (Fig. 2C). Since the analyzed group of treatment-naïve CRC patients was very homogenous with respect to pathological tumor (pT) status, pathological nodes (pN) and disease grade (Table S5), we could not correlate Vps37 staining intensity with clinical disease staging.

Overall, these results corroborated the finding of our bioinformatics analysis that the abundance of *VPS37B* is decreased at transcriptional and protein levels in CRC. They further mounted evidence for reduced mRNA and protein levels of *VPS37A* in CRC.

### Concurrent depletion of Vps37 proteins induces multifaceted transcriptional responses in CRC cells

Humans have four *VPS37* genes whose protein products display distinct domain architecture suggesting partly different functions in the cells. They all possess the Mod(r) domain mediating interaction with the remaining core ESCRT-I subunits. Whereas only Vps37A has the ubiquitin-binding UEV domain, the other members contain the proline-rich region (PRR) essential for protein-protein interactions (Fig. 3A). To study the cellular functions of Vps37 paralogs, we used an *in vitro* model of human colon cancer DLD1 cell line. The expression levels of *VPS37* paralogs in the DLD1 cell line reflect those observed in samples of CRC patients deposited in TCGA (Dou et al., 2016), as well in our data (GSE152195).

**Fig. 3.**
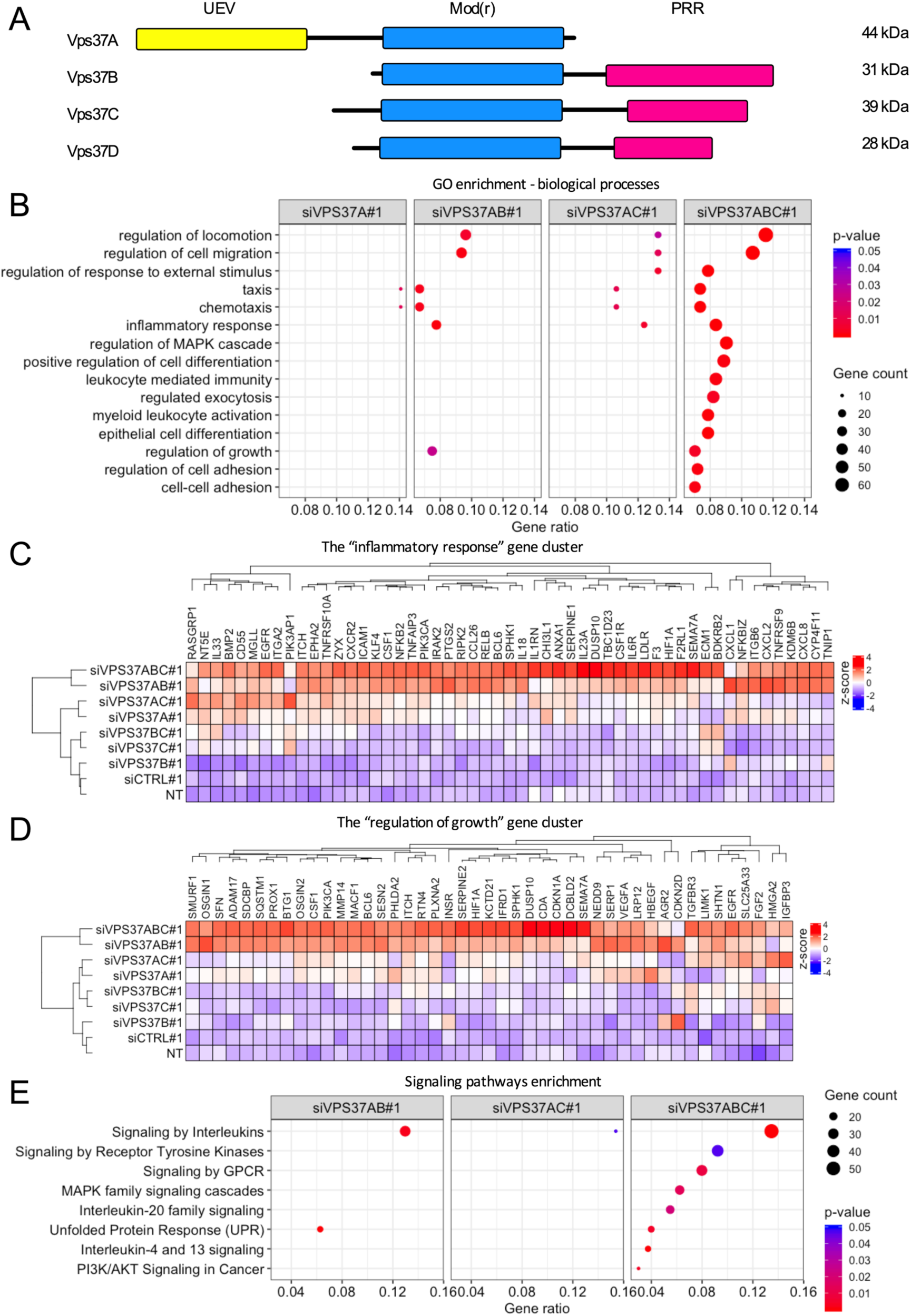
Concurrent depletion of Vps37 proteins induces multiple transcriptional responses in DLD1 cells. (A) Domain architecture of human *VPS37* paralogs. UEV – ubiquitin enzyme variant, Mod(r) – modifier of rudimentary, PRR – proline rich region. (B) Top 15 biological processes from gene ontology (GO) analysis among differentially expressed genes (≥1.50-fold or ≤0.667-fold; adjusted *P*<0.05) after individual or concurrent silencing of *VPS37* paralogs. Analysis was performed using the enrichGO function from clusterProfiler. (C-D) Heatmaps visualizing expression of genes related to inflammatory response (C) and regulation of cell growth (D) generated from the GO analysis of biological processes across different transfection conditions. (E) Selected pathways from the signaling pathway analysis among differentially expressed genes after individual or concurrent silencing of *VPS37* paralogs. Analysis was performed using the enrichPathway function from ReactomePA. RNA-Seq data analysis was performed using n=3 independent experiments. NT – non-transfected cells, abbreviations for differentially transfected cells are explained in Materials and Methods.

To gain insights into the molecular consequences of individual and concurrent depletion of Vps37 proteins on cellular homeostasis, we performed RNA-Seq in DLD1 cells. We first verified high knockdown efficiency and selectivity of siRNA (two independent sequences per target) by measuring the protein abundance of the three paralogs in DLD1 cells (Fig. S2A-C). Despite the sequence similarity between *VPS37A, VPS37B* and *VPS37C* (Fig. 3A), we could selectively silence the expression of each individual paralog or their combinations. As we observed certain differences in the silencing efficiency, we used single strongly acting siRNAs for RNA-Seq to knockdown *VPS37* paralogs individually or in double and triple combinations (7 conditions), compared to non-transfected cells (NT) and cells transfected with a combination of non-targeting siRNA (siCTRL#1). Nevertheless, all subsequent experiments were performed with two independent siRNA sequences per target. We considered genes to be differentially expressed when their expression was either below 0.667-fold or above 1.5-fold, and adjusted *P*<0.05 when normalizing against the two control conditions – siCTRL#1 and non-transfected cell (NT) (Fig. S2D).

We observed that co-silencing of *VPS37A, VPS37B* and *VPS37C* (abbreviated as *VPS37ABC*) elicited the greatest transcriptional changes (1277 genes). Pronounced changes (781 genes) were also detected after concurrent silencing of *VPS37AB* indicating the importance of these two subunits in maintaining homeostasis of DLD1 cells. Conversely, a limited number of genes underwent transcriptional changes upon other silencing combinations of *VPS37* family members (Fig. S2D). Hierarchical clustering of all investigated conditions on a set of differentially expressed genes after individual, double and triple silencing demonstrated that the branch containing siVPS37ABC#1 and siVPS37AB#1 was clearly distinct from the remaining conditions (Fig. S2E). We further observed that the pools of genes induced upon single depletion of Vps37A, Vps37B and Vps37C were largely non-overlapping (Fig. S2F), indicating that multifaceted transcriptional responses induced upon co-depletion of Vps37 proteins stem from the accumulation of paralog-specific defects in the cells.

Differentially expressed genes under each silencing conditions were subjected to the Gene Ontology (GO) analysis of biological processes. We identified biological processes only in transfection conditions with knockdown of the *VPS37A* paralog: siVPS37A#1, siVPS37AB#1, VPS37AC#1 and VPS37ABC#1. Among the top 15 gene signatures (whose order was determined based on a number of genes in the cluster upon *VPS37ABC* silencing) were processes related to cell migration, cellular signaling, inflammatory response, cell growth, and adhesion (Fig. 3B). We further focused on the “inflammatory response” (GO:0006954) and “regulation of growth” (GO:0040008) gene clusters. The inflammatory response heatmap contained genes encoding cytokines (*CXCL8*), adhesion molecules (*ICAM1*), and negative regulators of NF-κB signaling (*TNFAIP3*). In-depth interrogation of genes linked to the “regulation of growth” showed the presence of cyclin-dependent kinase inhibitors (*CDKN1A, CDKN2D*) and regulators of cell growth (*HMGA2, SFN*) (Fig. 3C,D). To determine the signaling pathways associated with inflammatory response and regulation of cell growth, we conducted a pathway network analysis using the Reactome Database. Our analysis of differentially expressed genes yielded enrichment of annotations related to signaling initiated by cytokines, receptor tyrosine kinases, and G-protein coupled response and involving mitogen-activated protein kinases (MAPK) and PI3K/Akt (Fig. 3E).

Collectively, these data point to largely non-redundant cellular functions of *VPS37* paralogs. The type and magnitude of transcriptional responses after their co-silencing are the cumulative response to perturbations of individual functions executed by Vps37 proteins. Concurrent knockdown of *VPS37AB* profoundly affects gene expression patterns linked to “inflammatory response” and “regulation of cell growth” and additional silencing of *VPS37C* paralog on top of *VPS37AB* knockdown further potentiates perturbations in gene transcription. These data suggested that Vps37 depletion activates multifaceted stress responses in the cells.

### Inflammatory gene expression is induced upon concurrent depletion of Vps37 proteins

To validate our RNA-Seq analysis, we selected the most pronouncedly induced genes (based on the fold change values) from to the inflammatory response cluster that represented different classes of molecules (cyto-/chemokine, adhesion molecule, classical and non-classical regulators of NF-κB signaling; Fig. 3C, GSE152195), and performed a qRT-PCR analysis of DLD1 cells subjected to individual or concurrent silencing of *VPS37* paralogs. We found that knockdown of *VPS37A* induced transcription of *TNFAIP3* (encoding A20) as well as *ICAM1* and had modest, yet insignificant, the effect on the expression of *CXCL8* (encoding IL-8) and *NFKBIA* (encoding IκBα) (Fig. S3A-D). Silencing of either *VPS37B* or *VPS37C* did not affect any of the investigated genes. Concurrent Vps37AB depletion significantly promoted *CXCL8, NFKBIA* and *TNFAIP3* transcription and modestly affected *ICAM1* transcription. Expression of the *TNFAIP3* gene was also induced by concurrent *VPS37AC* silencing. Finally, we observed that concurrent knockdown of *VPS37ABC* increased the expression of *CXCL8, ICAM1, NFKBIA*, and *TNFAIP3* comparable to concurrent Vps37AB depletion (Fig. S3A-D).

Since the magnitude of transcriptional changes in DLD1 cells was modest, we tested the expression of the same genes in a poorly differentiated RKO carcinoma cell line. We found that knockdown of *VPS37A* had a modest, yet insignificant, effect on the expression of *CXCL8* and *ICAM1*, whereas it did not affect *TNFAIP3* or *NFKBIA* transcription (Fig. 4A-D). Silencing of either *VPS37B* or *VPS37C* did not affect the expression of any of the investigated genes. Concurrent *VPS37AB* silencing induced an increase of *CXCL8, ICAM1, TNFAIP3*, and *NFKBIA* expression. Neither knockdown of *VPS37AC* nor *VPS37BC* substantially affected the transcription of the investigated targets (Fig. 4A-D). We observed that concurrent depletion of Vps37ABC had further positive effects on the magnitude of *CXCL8, ICAM1, NFKBIA* and *TNFAIP3* transcription compared to the Vps37AB-depleted cells (Figs 4A-D, S3E-G). Since the effects of individual and concurrent depletion of Vps37 proteins on gene expression were more pronounced in RKO cells, we decided to use it as the main model system in the subsequent experiments.

**Fig. 4.**
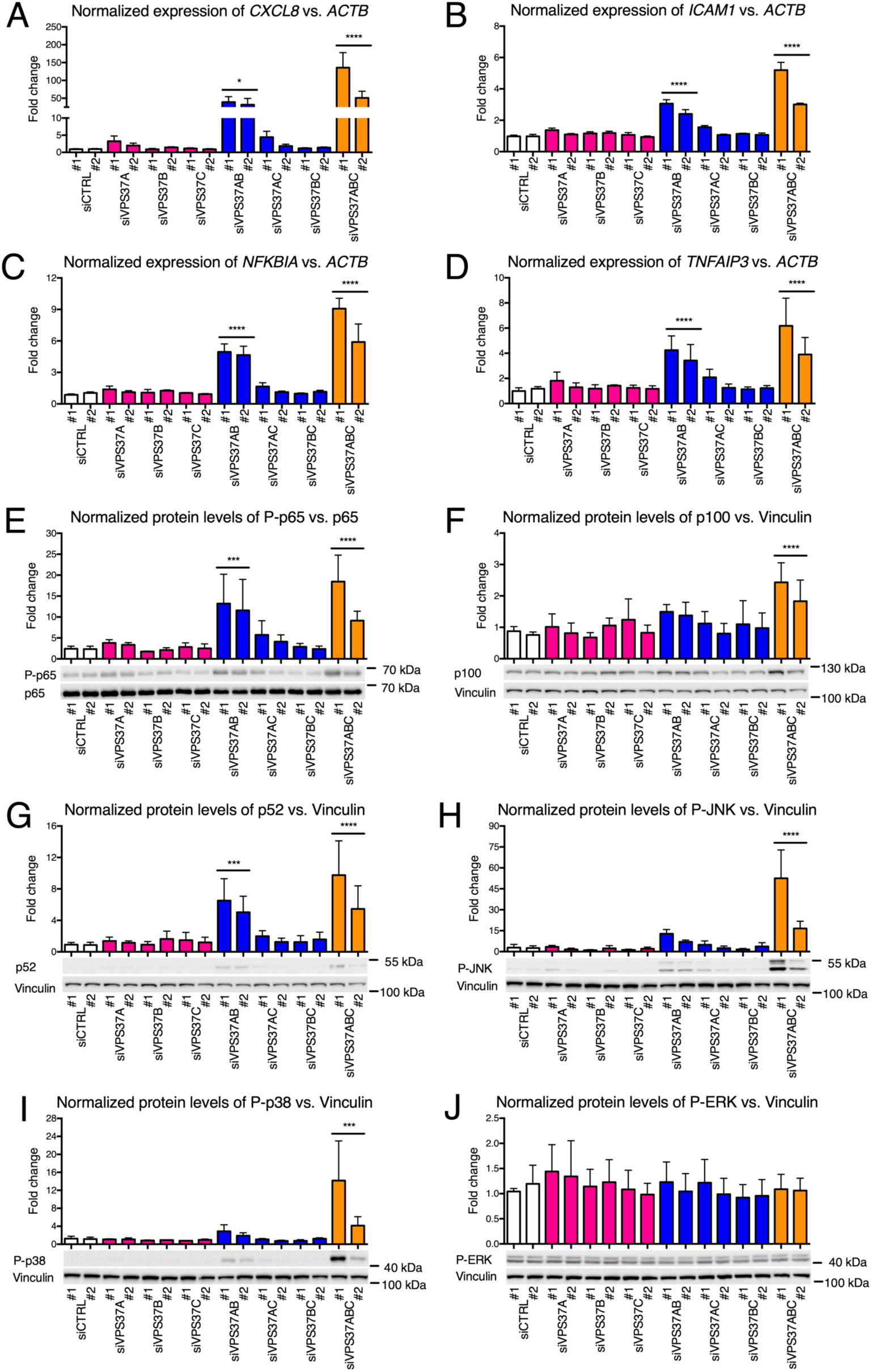
Concurrent depletion of Vps37 proteins induces MAPK and NF-κB inflammatory responses in RKO cells. (A-D) qRT-PCR analysis of expression of selected genes in the inflammatory response cluster: (A) *CXCL8*, (B) *ICAM1*, (C) *NFKBIA* and (D) *TNFAIP3*, measured 72 h after transfection with siRNA targeting *VPS37* paralogs individually or in combinations. Non-transfected (NT) cells and transfected with non-targeting siRNA (siCTRL) were used to assess the basal expression level of the investigated genes. *ACTB* (encoding β-Actin) was used as a reference gene. (E-G) Western blotting analysis of the NF-κB pathway activation: (E) phosphorylation of p65, (F) p100 and (G) p52 protein abundance. (H-J) Western blotting analysis of MAPK activation: (H) phosphorylation of JNK, (I) phosphorylation of p38 and (J) phosphorylation of ERK. (E-J) Lysates of RKO cells were collected 72 h after transfection with siRNA targeting *VPS37* paralogs individually or in combinations. Lysates from non-transfected (NT) cells and transfected with non-targeting siRNA (siCTRL) were used to assess the basal level of intracellular signaling. p65 and vinculin were used as loading controls. Representative blots are shown along with densitometry analysis. Data in all panels are mean ± standard deviation of n=3 independent experiments expressed as the fold change of either mRNA (A-D) or protein (E-J) levels in NT cells, which was set as 1. Statistical significance was assessed against grouped siCTRL conditions using one-way ANOVA test followed by Bonferroni’s correction; \**P*<0.05, \*\**P*<0.01, \*\*\**P*<0.001, \*\*\*\**P*<0.0001.

In summary, we found that concurrent depletion of Vps37 proteins, in particular Vps37AB and Vps37ABC, activates transcription of multiple classes of inflammatory genes, while knockdown of individual *VPS37* paralogs induces only minor changes in their expression. We also noted that the rate of gene transcription is cell-type dependent.

### Concurrent knockdown of *VPS37* paralogs induces NF-κB signaling and MAPK cascade

We and others previously revealed that depletion of the core ESCRT-I subunits – Tsg101 and Vps28 – induced NF-κB-driven inflammatory response (Brankatschk et al., 2012; Maminska et al., 2016). As our RNA-Seq analysis yielded the “inflammatory response” gene cluster upon siVPS37AB#1 and siVPS37ABC#1 (Fig. 3C), we combined genes from each cluster into a single list and subjected them for transcriptional motif-enrichment analysis using RcisTarget (Aibar et al., 2017). For each gene, the promoter region of 500 bp upstream and 100 bp downstream to the transcription starting site was investigated with the transcription factor binding sites (TFBS) matrices available in the JASPAR database. Our *in silico* analysis revealed enrichment of TFBS for the members of NF-κB, FOS, and AP1 transcription factors. The consensus sequence for RELA/p65 with the highest normalized enrichment score was the top annotated TFBS (Table 2).

Because concurrent silencing of *VPS37* paralogs enhanced the expression of NF-κB target genes (Figs 4A-D, S3A-D, Table 2), we explored the molecular basis of these effects. To this end we measured p65 phosphorylation and p100 to p52 processing as hallmarks of canonical and non-canonical NF-κB signaling, respectively. Our Western blot analysis of lysates from RKO cells with silencing of individual *VPS37* paralogs as well as *VPS37AC* and *VPS37BC* showed no significant induction of any branch of the NF-κB pathway (Fig. 4E-G). *VPS37AB* knockdown induced p65 phosphorylation and cleavage of p100 to p52. Co-silencing of *VPS37ABC* induced p65 phosphorylation, cleavage of p100 to p52 (Fig. 4E, G), and compared to Vps37AB depletion also increased p100 levels (Fig. 4F).

MAPKs cooperate with NF-κB in driving inflammation (Hoesel and Schmid, 2013). Since signaling enrichment analysis of our transcriptomics data pointed to increased expression of genes whose products regulate the MAPK cascade, we tested phosphorylation of JNK, p38 and ERK as hallmarks of MAPK activation. Using Western blotting, we found that silencing of individual *VPS37* paralogs did not affect JNK, p38, and ERK phosphorylation (Fig. 4H-J). Knockdown of *VPS37AB* in RKO cells had minor positive, yet insignificant, effect on phosphorylation of JNK and p38 MAPK but did not activate ERK. Neither *VPS37AC* nor *VPS37BC* silencing activated JNK, p38 and ERK (Fig. 4H-J). Concurrent Vps37ABC depletion induced JNK and p38 phosphorylation but again it did not activate ERK (Fig. 4H-J).

Overall, we concluded that depletion of Vps37ABC is associated with the activation of canonical and non-canonical NF-κB signaling as well as JNK and p38 MAPK.

### Cell proliferation and colony forming ability of CRC cells are inhibited after concurrent knockdown of *VPS37* paralogs

As GO analysis of biological processes revealed the regulation of growth gene cluster (Fig. 3D, GSE152195), we examined the effect of differential depletion of Vps37 proteins on cell growth using a short-term BrdU proliferation assay and a long-term colony formation assay in DLD1 and RKO cells.

Using the proliferation assay, we observed that single silencing of *VPS37A* modestly decreased the proliferation rate, which was statistically significant only in RKO cells (Fig. 5A,B). Knockdown of neither *VPS37B* nor *VPS37C* altered growth of DLD1 and RKO cells. Silencing of *VPS37AB* significantly inhibited DLD1 and RKO cell proliferation, whilst knockdown of *VPS37AC* and *VPS37BC* had no impact. The strongest inhibition of cell proliferation was seen upon *VPS37ABC* knockdown (Fig. 5A,B). In line with the results of proliferation assay, depletion of Vps37A alone or co-silencing of *VPS37AB, VPS37AC* and *VPS37ABC* inhibited ability of DLD1 and RKO cells to form colonies in the clonogenic assay performed 14 days after siRNA transfection (Figs 5C-D, S4A-B). In parallel, we checked the impact of *TSG101* silencing on cell proliferation and colony formation. Its knockdown in RKO cells inhibited both processes, comparably to concurrent silencing of *VPS37* paralogs (Fig. S4C-E).

**Fig. 5.**
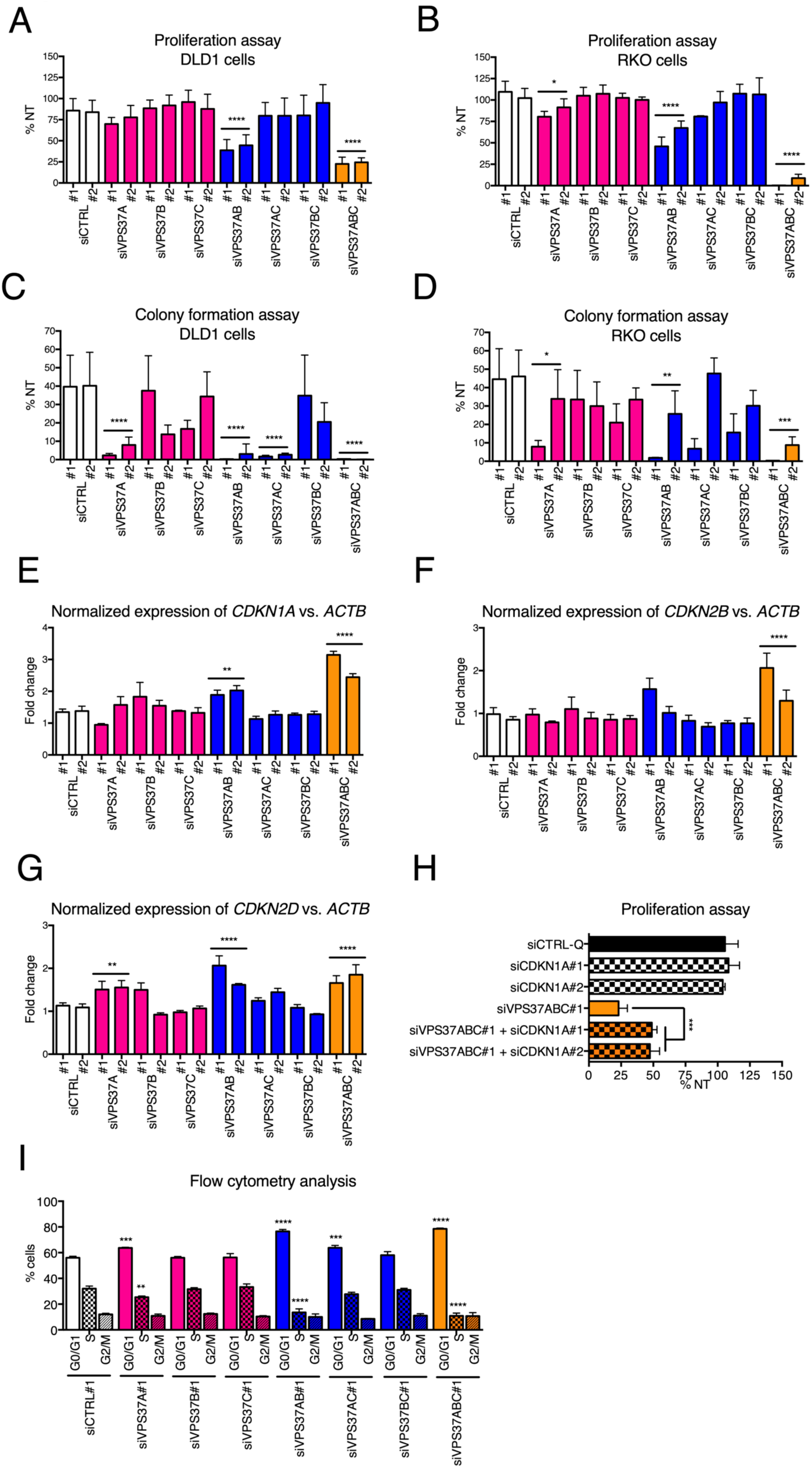
Concurrent depletion of Vps37 proteins inhibits CRC cell growth *in vitro*. (A, B) Cell proliferation of (A) DLD1 and (B) RKO cells assessed 120 h after individual and concurrent depletion of *VPS37* paralogs using the BrdU proliferation assay. (C, D) Clonogenic growth of (C) DLD1 and (D) RKO cells assessed 14 days after individual and concurrent knockdown of *VPS37* paralogs with representative images for each transfection condition shown in Fig. S4A for DLD1 cells and Fig. S4B for RKO cells. (E-G) qRT-PCR analysis of expression of genes encoding *CDKNs*: (E) *CDKN1A*, (F) *CDKN2B* and (G) *CDKN2D*, measured 72 h after individual and concurrent depletion of Vps37 paralogs. Non-transfected (NT) cells and transfected with non-targeting siRNA were used to assess the basal level of the investigated genes. *ACTB* (encoding β-Actin) was used as a reference gene. (H) Cell proliferation of RKO cells was assessed 120 h after concurrent silencing of all *VPS37* paralogs and *CDKN1A* using the proliferation assay. (I) Analysis of cell cycle was performed upon individual or concurrent silencing of *VPS37* paralogs. Cells were forward transfected for 96 h, stained with PI and evaluated with a flow cytometer. Data presented in all panels are mean of n=3 independent experiments ± standard deviation analyzed with one-way ANOVA with Bonferroni’s correction. Statistical significance for grouped siCTRL conditions (A-G, I) and for siCTRL#1-Q (I); \**P*<0.05; \*\**P*< 0.01, \*\*\**P*< 0.001, \*\*\*\**P*< 0.0001.

In conclusion, we found that concurrent depletion of Vps37 proteins has detrimental effects on cancer cell growth *in vitro* and the phenotype of Tsg101-depleted cells closely resembles the one observed in *VPS37ABC* knockdown cells. Our results also indicate that growth rate is primarily dependent on the expression of both *VPS37A* and *VPS37B*. Additional co-silencing of *VPS37C* potentiated proliferation and colony-forming defects of *VPS37AB* knockdown. In contrast, long-term growth rate appears to be largely dependent on Vps37A.

### Concurrent depletion of Vps37 proteins induces p21-mediated cell cycle arrest

Accelerated division of tumor cells is among others a result of abnormal activity of cyclins and cyclin-dependent kinase inhibitors (CDKNs) (Bonelli et al., 2014). As our RNA-Seq data revealed increased expression of three genes encoding *CDKNs* (*CDKN1A, CDKN2B*, and *CDKN2D*, GSE152195, Fig. 3D), we used qRT-PCR to corroborate their changed transcription in RKO and DLD1 cells subjected to individual and concurrent silencing of *VPS37* paralogs. In RKO cells, knockdown of *VPS37A* did not affect *CDKN1A* or *CDKN2B* transcription (Fig. 5E-F) but modestly induced the expression of *CDKN2D* (Fig. 5G). Silencing of either *VPS37B* or *VPS37C* did not change transcription of any of the analyzed genes. We found that *VPS37AB* silencing increased *CDKN1A, CDKN2B*, and *CDKN2D* transcription; yet in the case of *CDKN2B*, the increase was observed for only one pair of siRNA and did not reach statistical significance. Knockdown of neither *VPS37AC* nor *VPS37BC* paralogs had an impact on the investigated target genes (Fig. 5E-G). Instead, concurrent depletion of Vps37ABC further increased transcription of all investigated *CDKNs* compared to concurrent depletion of Vps37AB (Fig. 5E). In line, we found a similar pattern of *CDKN* expression in differentially transfected DLD1 cells; however, we could not corroborate enhanced *CDKN2D* transcription upon co-depletion of Vps37 proteins that we initially identified in our RNA-Seq analysis (Fig. S4F-H). Finally, we observed that the transcription pattern of *CDKNs* after depletion of Tsg101 in RKO cells paralleled those observed for *CDKN1A* and *CDKN2B* expression after concurrent *VPS37ABC* silencing. In this case, silencing of *TSG101* did not induce *CDKN2D* expression (Fig. S4I).

Increased expression of *CDKNs* after concurrent Vps37ABC depletion suggests an impact on the proliferation rate and cell cycle progression. *CDKN1A* encodes p21, which inhibits cell cycle progression in the G1, S and G2 phases, whilst *CDKN2B* and *CDKN2D* encode p15^INK4B^ and p19^INK4D^, respectively, which inhibit complexes formed by cyclin D and halt cell cycle in the G1 phase (Bonelli et al., 2014). Thus, we evaluated the proliferation of Vps37ABC-depleted RKO cells upon co-silencing of the *CDKN1A* gene, which was the most pronouncedly induced in our qRT-PCR analysis. We observed that concurrent knockdown of *VPS37ABC* and *CDKN1A* partly rescued cell proliferation, corroborating the inhibitory impact of p15^INK4B^ and p19^INK4D^ on cell division (Fig. 5H). In line, p21 depletion in cells transfected with siRNA against *TSG101* improved RKO cell proliferation (Fig. S4J).

To gain further insights into the inhibition of cell growth after differential silencing of *VPS37* paralogs, siRNA-transfected RKO cells were stained with propidium iodide and cell cycle was analyzed by flow cytometry. We observed that knockdown of either *VPS37B* or *VPS37C* did not change cell cycle progression as indicated by the unaltered percentage of cells in the G0/G1 and S phases (Fig. 5I). Knockdown of *VPS37A* increased the percentage of cells in the G0/G1 phase and decreased the number of cells in the S phase (Fig. 5I). The impact of Vps37AC-depletion closely paralleled that observed after *VPS37A* silencing. In contrast to Vps37BC-depleted cells, whose cell cycle progression was not affected, concurrent knockdown of *VPS37AB* resulted in the increased number of cells in the G0/G1 phase and a drop in the S phase (Fig. 5I). The proportion of cells in the G0/G1 phase after combined silencing of *VPS37ABC* was comparable to that observed in Vps37AB-depleted cells (Fig. 5I). Finally, silencing of *TSG101* closely paralleled the effects observed after *VPS37ABC* knockdown (Fig. S4K). None of the analyzed silencing conditions (involving *VPS37* paralogs and *TSG101*) altered the percentage of cells in the G2/M phase (Figs 5I, S4K).

In summary, our data uncovered that concurrent depletion of Vps37 proteins induces the expression of three *CDKNs*, which cooperatively halt cell cycle in the G1 phase. Moreover, the phenotype of Tsg101-depleted cells closely resembles the one observed in *VPS37ABC* knockdown cells. We further demonstrated that *CDKN* expression and cell cycle progression are primarily dependent on Vps37A and modulated by the presence of other family members.

### NF-κB response and p21-mediated growth arrest are induced independently after depletion of *VPS37* paralogs

We next investigated the molecular basis for the induction of NF-κB response and p21-mediated growth arrest after depletion of all three Vps37 proteins in RKO cells. Since *CDKN1A* encoding p21 was the most potently affected gene in our qRT-PCR analysis (Fig. 5E) and its knockdown in Vps37ABC-depleted RKO cells partly rescued their proliferation (Fig. 5H), we used it as readout to assess the relationship between inflammatory response and cell growth arrest in Vps37ABC-depleted cells.

We first investigated the time course of changes in p21 levels and activation of the NF-κB pathway components after concurrent *VPS37ABC* silencing in RKO cells. We observed that depletion of Vps37ABC increased p21 abundance after 24 h and 72 h post-transfection (Fig. 6A). We found rapid phosphorylation of p65 in Vps37ABC-depleted cells, 24 h and 72 h post-transfection; however, after 24 h p65 phosphorylation did not reach statistical significance (Fig. 6B). The abundance of p100/p52 increased from 24 h to 72 h post-transfection in cells with *VPS37ABC* knockdown (Fig. 6C,D). Throughout 24-72 h post-transfection, abundance of Vps37A, Vps37B and Vps37C gradually decreased after 24 h and remained undetectable after 72 h post-transfection (Fig. S5A-C). These data showed that the activation of NF-κB signaling and production of p21 occurred within the similar timeframe after Vps37ABC depletion, thus none of these processes preceded each other.

**Fig. 6.**
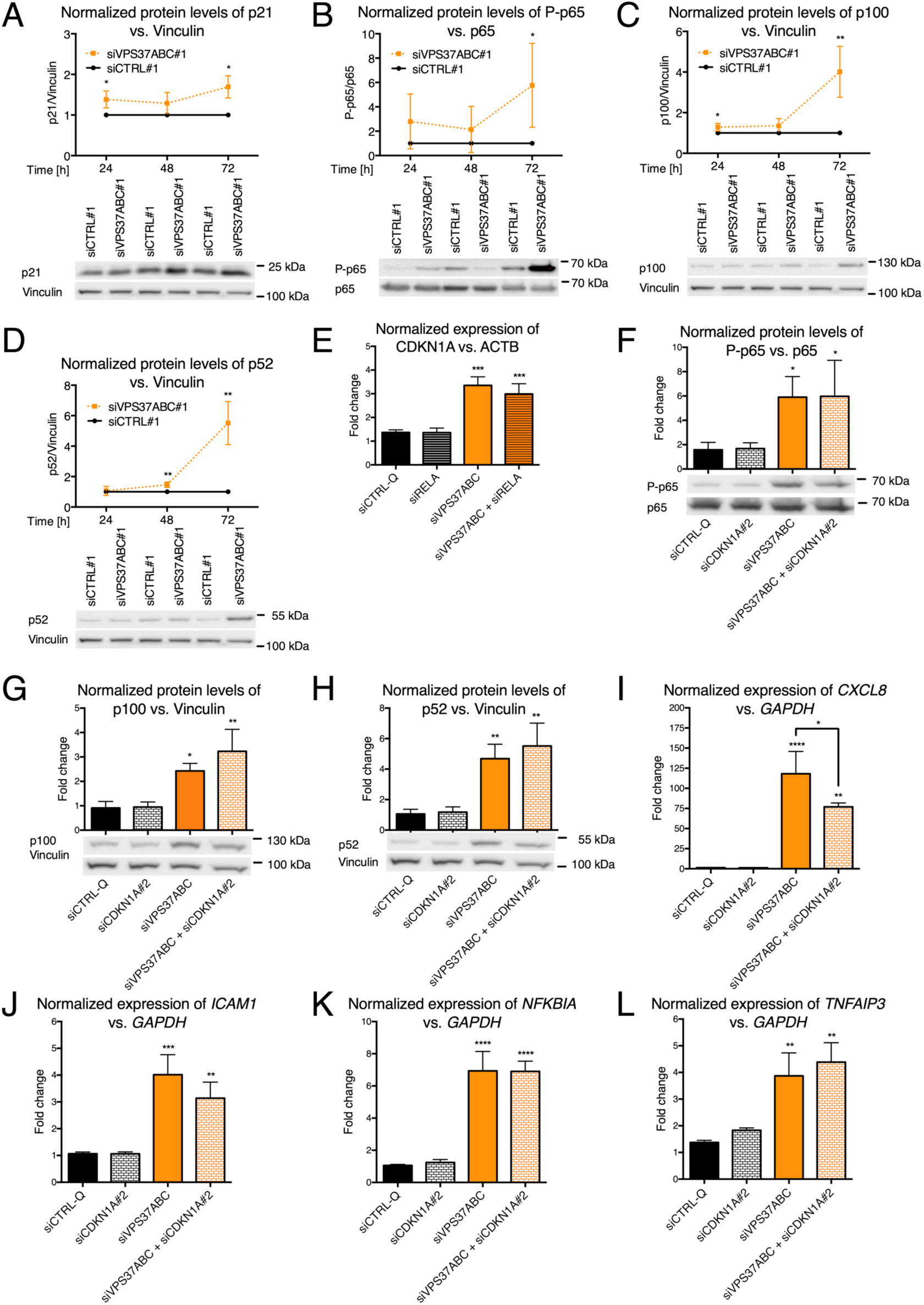
Inflammatory response and cell growth inhibition after concurrent depletion of *VPS37* paralogs are two independently regulated processes in RKO cells. (A-D) Western blot analyses of (A) p21, (B) phosphorylated p65, (C) p100 and (D) p52 abundance were performed 24, 48 and 72 h after transfection with non-targeting or on-target siRNA for *VPS37ABC*. (E) qRT-PCR analysis of *CDKN1A* was performed 72 h after transfection with siRNA targeting the p65 subunit of NF-κB dimer (*RELA*) alone or in combination with three *VPS37* paralogs. (F-H) Western blotting analyses of (F) phosphorylated p65, (G) p100 and (H) p52 abundance were performed 72 h after transfection with siRNA targeting CDKN1A alone or in combination with all the three *VPS37* paralogs. (I-L) qRT-PCR analyses of (I) *CXCL8*, (J) *ICAM1*, (K) *NFKBIA*, and (L) *TNFAIP3* were performed 72 h after transfection with siRNA targeting *CDKN1A* alone or in combination with all the three *VPS37* paralogs. (E, I-L) *ACTB* (encoding β-Actin) or *GAPDH* (encoding Glyceraldehyde 3-phosphate dehydrogenase) were used as a reference gene in qRT-PCR analysis. (A-D, F-H) p65 and vinculin were used as loading controls for Western blotting. Representative blots are shown along with densitometry analysis. Data in all panels are mean ± standard deviation of n=3 independent experiments expressed as the fold change of either mRNA or protein level. In panels A-D protein abundance in siCTRL#1 transfected cells was set to 1, whilst in panels E-L mRNA and protein abundance in non-transfected (NT) cells was set as 1. Statistical significance was assessed using unpaired Student’s t-test (A-D) or one-way ANOVA test followed by Bonferroni’s correction (E-L). Statistical significance against siCTRL#1 at matching time point for transfection with three different siRNA (A-D) and siCTRL-Q for four different siRNA (E-L), \**P*<0.05, \*\**P*<0.01, \*\*\**P*<0.001, \*\*\*\**P*<0.0001.

In certain cell types, the canonical NF-κB pathway is crucial for *CDKN1A* transcription (Ledoux and Perkins, 2014) and we verified whether increased *CDKN1A* expression after co-depletion of Vps37 proteins required the canonical NF-κB subunit p65 (encoded by *RELA*). Knockdown of *RELA* in Vps37ABC-depleted cells did not affect *CDKN1A* expression (Fig. 6E), although it almost completely blunted transcription of *CXCL8* (Fig. S5D) and substantially reduced expression of *ICAM1*, two prototypical NF-κB target genes (Figs S5E, S5F-I). These results suggest that negative effects on cell growth stemming from co-silencing of *VPS37* paralogs are not consequences of induction of canonical NF-κB signaling.

p21 modules NF-κB signaling in immune cells (Rackov et al., 2016; Trakala et al., 2009) but whether similar mechanisms occur in CRC cells has not been assessed. Thus, we explored whether silencing of *CDKN1A* affected phosphorylation of p65 and processing of p100 to p52 upon Vps37ABC depletion. As assessed by Western blot, co-silencing of *VPS37* paralogs and *CDKN1A* did not affect the levels of p65 phosphorylation and p100 to p52 processing compared to co-silencing of *VPS37* paralogs alone (Figs 6F-H, S5J-M). We also checked whether p21 depletion modulated the rate of inflammatory gene expression in Vps37ABC-depleted cells. Co-silencing of *CDKN1A* inhibited transcription of only *CXCL8* but not *ICAM1, TNFAIP3* and *NFKBIA* (Fig. 6I-L). These results showed no modulatory impact of p21 on NF-κB signaling and three out of four investigated target genes.

Overall, we concluded that in CRC cells with co-depletion of Vps37 proteins the induction of NF-κB inflammatory response and p21-mediated cell growth inhibition are two independent processes. These data point out that cell growth arrest is not caused by activation of inflammatory response.

### ESCRT-I is destabilized after either concurrent depletion of Vps37 proteins or *TSG101* silencing

We speculated that the type and magnitude of transcriptional responses after individual and concurrent silencing of *VPS37* paralogs might be attributed to distinct ESCRT-I stability. It was previously shown that knockdown of some ESCRT-I core components induced partial or complete degradation of other complex subunits (Stefani et al., 2011; Wunderley et al., 2014); yet, a detailed characterization of all ESCRT-I subunits after individual and concurrent Vps37 proteins depletion has not been performed so far.

First, we checked whether knockdown of individual ESCRT-I components affected the stability of its remaining subunits expressed in CRC cells. Western blot analysis of lysates from RKO cells revealed that depletion of either Tsg101 or Vps28 destabilized each other (Fig. S6A,B) as well as Vps37A, Vps37B, Vps37C, Mvb12A, Mvb12B, and lowered UBAP-1 protein abundance (Fig. S6C-H). We observed that silencing of *VPS37A* diminished UBAP-1 protein abundance indicating that ESCRT-I complexes containing Vps37A preferentially incorporate UBAP-1 (Fig. S6F). Depletion of Vps37B reduced the abundance of Tsg101 protein (Fig. S6A) and partially Mvb12A (Fig. S6G). Conversely, silencing of *MVB12A* decreased Vps37B (Fig. S6D), indicating partnering preference between these subunits. We did not observe any relationship between the stability of Vps37C and Mvb12 proteins (Figs S6E, S6G-H).

We next analyzed the stability of ESCRT-I core and auxiliary subunits upon concurrent silencing of two or three *VPS37* paralogs. Co-depletion of Vps37AB or Vps37BC proteins decreased Tsg101 and Vps28 abundance, whilst the effects of *VPS37AC* knockdown were less potent (Fig. 7D,E). Knockdown of all three *VPS37* genes led to the complete destabilization of Tsg101 and Vps28 proteins (Fig. 7D-E). Protein abundance of UBAP-1 was decreased in all silencing combinations involving *VPS37A* (Fig. 7F) corroborating the results of individual *VPS37A* knockdown (Fig. S6F). Similarly, Mvb12A abundance was reduced whenever cells were depleted of Vps37B (Fig. 7G), whilst such effect was less pronounced for Mvb12B (Fig. 7H). Silencing of all *VPS37* genes depleted both Mvb12 proteins (Fig. 7G,H). Notably, the stability of the core and auxiliary ESCRT-I subunits after concurrent *VPS37ABC* knockdown closely resembled the effects of *TSG101* silencing (Fig. S6A).

**Fig. 7.**
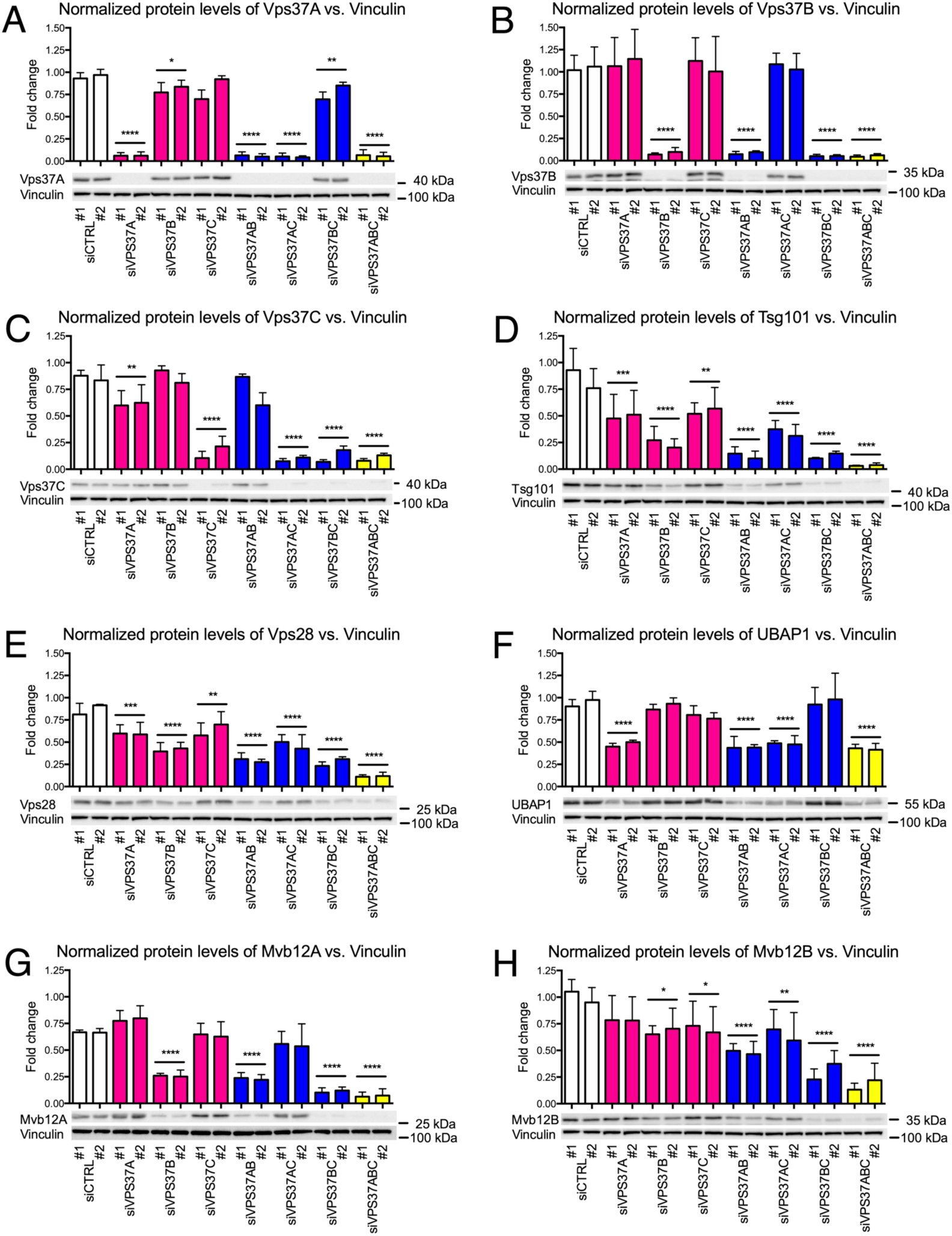
Concurrent depletion of Vps37 proteins destabilizes ESCRT-I components and reveals partnering preferences for auxiliary subunits. Western blotting analysis of ESCRT-I subunits: (A) Vps37A, (B) Vps37B, (C) Vps37C, (D) Tsg101, (E) Vps28, (F) UBAP-1, (G) Mvb12A and (H) Mvb12B. Lysates of RKO cells were collected 72 h after transfection with siRNA targeting *VPS37* paralogs individually or in combinations. Lysates of RKO cells from non-transfected (NT) cells and transfected with non-targeting siRNA were used to assess the basal level of ESCRT-I subunits. Vinculin was used as a loading control. Representative blots are shown along with densitometry analysis. Data are mean ± standard deviation of n=3 independent experiments expressed as the fold change of protein level in NT cells, which was set as 1. Statistical significance was assessed using ANOVA test followed by Bonferroni’s correction. Statistical significance against siCTRL conditions, \**P*<0.05, \*\**P*<0.01, \*\*\**P*<0.001, \*\*\*\**P*<0.0001.

In summary, these data show an inter-dependability of ESCRT-I subunits for maintaining the complex stability. They indicate that the incorporation of auxiliary subunits is selective with respect to their Vps37 partners (Vps37A with UBAP-1 and Vps37B with Mvb12A). We also found that simultaneous interference with *VPS37A* and *VPS37B* expression induces pronounced decrease in ESCRT-I stability and Vps37C depletion only slightly magnifies this effect. Our results further argue that the type and magnitude of transcriptional responses after differential depletion of Vps37 proteins correlate with the abundance of core and accessory ESCRT-I components.

## DISCUSSION

Tumors develop various mechanisms to prolong exposure of plasma membrane receptors to ensure constitutive signaling that is beneficial for their growth. One of such mechanisms relies on altered expression of endocytic transport regulators (Barbieri et al., 2016; Floyd and De Camilli, 1998; Mellman and Yarden, 2013; Mosesson et al., 2008; Schmid, 2017). The advent of next-generation sequencing technologies has permitted unbiased screening of components orchestrating receptor transport and endo-lysosomal degradation in distinct pathological settings (Buser et al., 2019; Yoshida et al., 2010). Here, by mining TCGA data we revealed differential expression of 113 endocytic machinery components across stages of CRC, several of which were previously shown to be reduced in adenocarcinoma (Kwong et al., 2005; Szymanska et al., 2020; Tanigawa et al., 2019). In-depth validation of our *in silico* screen uncovered reduced mRNA and protein abundance of *VPS37A* and *VPS37B* paralogs in CRC patients. Our *in vitro* data further allowed us to propose that Vps37 proteins have partly non-overlapping functions in the cell. We also showed that co-depletion of Vps37 proteins evokes stress responses manifested among others by activation of the NF-κB inflammatory response and p21-mediated impairment of cell growth. We finally correlated the magnitude of stress responses with the degree of ESCRT-I subunit destabilization after (co-)depletion of Vps37 proteins. While decreased abundance of *VPS37A* and/or *VPS37B* mRNA and proteins appears not to be an oncogenic driver per se, these passenger alterations might represent potential vulnerabilities of cancer cells to therapeutic treatment.

In humans, there are four *VPS37* paralogs with distinct chromosomal localization, number of splicing isoforms, and protein sequence identity. *VPS37A, VPS37B, VPS37C* and *VPS37D* are localized on chromosome 8p, 12q, 11q, and 7q, respectively. This different chromosomal localization of *VPS37* genes could favor independent regulatory mechanisms of expression in patho-physiological circumstances. Another layer of complexity is added by the incorporation of Vps37 proteins into ESCRT-I to form functionally distinct complexes in the cell (Stefani et al., 2011; Wunderley et al., 2014). Changes in mRNA and protein levels of *VPS37A* were previously documented in various cancer types, including liver, prostate, breast, ovarian, renal, lung, glioma, gastric, oral and oropharyngeal squamous cell carcinoma and colon cancer (Chen et al., 2015; Chen et al., 2020; Chen et al., 2018; Du et al., 2016; Fu et al., 2018; Lai et al., 2009; Perisanidis et al., 2013; Sun et al., 2017; Vasaikar et al., 2019; Wittinger et al., 2011; Wu et al., 2019; Xu et al., 2017a; Xu et al., 2017b; Xu et al., 2014; Xu et al., 2017c; Xu et al., 2003; Yang et al., 2016; Yang et al., 2017; Zhu et al., 2015). Several of these studies suggested that *VPS37A* acts as a tumor suppressor and its loss can serve as an adverse prognostic factor. However, the abundance of the remaining *VPS37* paralogs at mRNA and protein levels as well as their contribution to oncogenesis have not been investigated across cancer types and disease stages.

By mining the TCGA expression data, we revealed decreased expression of *VPS37B* during the transition from early to advanced stages of colorectal adenocarcinoma. Analysis of an independent cohort of patients corroborated results of our *in silico* prediction that *VPS37B* expression is indeed reduced in adenocarcinoma patients. Additionally, we found decreased expression of *VPS37A* in the analyzed group of patients. We further confirmed the lower abundance of both proteins in treatment-naïve primary CRC samples belonging largely to the locally advanced disease. The observations we made on *VPS37* paralog expression extend previous bioinformatics analysis of COAD and READ cohorts performed without grouping these patients based on disease stages (Miller et al., 2018) and a proteogenomic study of a homogenous cohort of treatment-naïve patients undergoing primary surgery for colon adenocarcinoma (Vasaikar et al., 2019). The *VPS37A* gene undergoes frequent deletion as a part of 8p during progression from adenoma to adenocarcinoma, which would explain its decreased abundance at mRNA and protein levels (Meijer et al., 1998). On the other hand, further studies will need to clarify the molecular origin of diminished *VPS37B* expression in CRC as chromosome 12q, where it is located, undergoes frequent amplifications, suggesting an existence of a (post-)transcriptional mechanism (Wood et al., 2007).

The present study provided comprehensive understanding of the consequences of Vps37 protein depletion in CRC cells in all possible combinations of paralogs expressed at high levels in these cells, namely *VPS37A, VPS37B* and *VPS37C*. Analysis of the transcriptome of CRC cells revealed that Vps37 proteins have partly non-overlapping function as deduced from distinct sets of genes induced upon their individual depletion. It also suggests that the type and magnitude of transcriptional responses upon concurrent *VPS37* paralog silencing stem from the cumulative inhibition of cellular processes executed by their protein products. Among genes induced upon either *VPS37AB* or *VPS37ABC* knockdown we did not find prototypical drivers of tumorigenesis but rather cell growth inhibitors, such as *CDKN1A*/p21, *CDKN2B*/p14^INK4B^, and *CDKN2D*/p19^INK4D^. As a consequence we observed decreased ability of CRC cells to progress over cell cycle phases that resulted in the inhibition of proliferation and colony forming ability. Our results from Vps37ABC-depleted cells reinforce the notion that knockdown of other ESCRT-I components, such as *TSG101, VPS28* and *UBAP1*, induces cell cycle arrest and halts cell proliferation (Krempler et al., 2002; Miller et al., 2018; Morita et al., 2007). Here, we also demonstrate that the degree of cell growth impairment depends primarily on the perturbed expression of *VPS37A* and whether it is silenced on its own or in combination with other paralogs. To our knowledge, expression of neither *VPS37B* nor *VPS37C* has been related to cancer growth but rather to virus release and infectivity (Stuchell et al., 2004). A vast majority of CRC cell lines tested within the DepMap project (Behan et al., 2019) showed no or only slight changes in cell fitness upon RNAi-mediated depletion of *VPS37A* or *VPS37B* (*VPS37C* has not been tested with this technology). Noteworthy, CRISPR-Cas9-mediated knockout of *VPS37A* markedly decreased RKO and DLD1 cell fitness, whilst the effects of *VPS37B* and *VPS37C* were less deleterious. Observations made within the DepMap project support the results of our colony formation and proliferation assays and point to distinct effects of long-and short-term loss of *VPS37* paralogs. Although our data suggest the negative impact of Vps37 paralogs (co-)depletion on cancer cell growth, we postulate that the effect of their CRC-associated reduction is more nuanced. Previous studies revealed that loss of chromosome 8p, where *VPS37A* is located, promotes tumor growth (Cai et al., 2016; Xue et al., 2012). Decreased expression of *VPS37* paralogs may be particularly beneficial for cells of advanced stage CRC as RNAi-mediated silencing of *VPS37A* promoted the resistance of prostate and breast cancer cells to chemotherapeutics (Sun et al., 2017; Yang et al., 2016). Our interrogation of TCGA expression data did not reveal increased transcription of *CDKN1A, CDKN2B* and *CDKN2D* in the advanced stages of CRC patients. Notably, this cohort displayed only reduced expression of *VPS37B* but not *VPS37A*, which at least in part might explain lack of changes in *CDKN* expression (Table S6). Thus, we postulate that the impact of *VPS37* paralog loss on cancer cell growth warrants further studies in a subset of patients with decreased expression of both *VPS37A* and *VPS37B*.

The pronounced inhibition of cell growth *in vitro* upon Vps37AB and Vps37ABC depletion correlated with activation of inflammatory and stress signaling mediated by NF-κB and MAPK. These cellular responses induced upon silencing of *VPS37* paralogs can be viewed as another example of sterile inflammation which resembles stress reactions caused by intracellular dysfunction of numerous membrane organelles, such as malfunctioning mitochondria, ER or endosomes (Keestra-Gounder et al., 2016; Maminska et al., 2016; West et al., 2015). Although Vps37B depletion did not promote inflammatory gene transcription in CRC cell lines, the subset of advanced stage CRC patients in the TCGA dataset with decreased *VPS37B* expression showed elevated mRNA abundance of *CXCL8* and *ICAM1* (Table S6). In our *in vitro* experimental setting, expression of these genes was increased only upon concurrent knockdown of either *VPS37AB* or *VPS37ABC*. If a subgroup of CRC patients with loss of both *VPS37A* and *VPS37B* was identified, it would be worth testing whether they display an inflammatory phenotype that could be modulated pharmacologically. However, inflammatory gene expression in advanced stage CRC patients is likely a result of multiple lesions accumulated in the course of disease progression. In addition to the expression of several NF-κB-dependent cytokines that clustered to processes related to (chemo-)taxis of immune system cells, *VPS37AB* and *VPS37ABC* knockdown cells produced high levels of the cell cycle inhibitor p21. Although several papers described NF-κB as a regulator of *CDKN1A*/p21 expression leading to cell cycle arrest in normal and cancer cells (Basile et al., 2003; Hinata et al., 2003; Nicolae et al., 2018; Wuerzberger-Davis et al., 2005), our data point to a different mechanism. Plausibly, it involves the release of Tsg101-mediated repression of the *CDKN1A* promoter (Lin et al., 2013), which in our study correlated with ESCRT-I destabilization after concurrent depletion of Vps37 proteins. Alternatively, ESCRT-I destabilization upon co-depletion of Vps37 proteins might induce p53-driven transcription of *CDKN1A* (El-Deiry, 2003). On the other hand, the p21 protein was shown to regulate the NF-κB pathway in macrophages (Rackov et al., 2016; Trakala et al., 2009), but we excluded that a similar mechanism occurs in CRC cells. Thus, we concluded that inflammatory response induction and inhibition of cell growth after concurrent *VPS37* paralog silencing are two independent and parallel processes.

The most important finding of this study is that differential depletion of Vps37 proteins elicits distinct effects on ESCRT-I subunit stability which align with the type and magnitude of transcriptional responses. Our data reinforce the notion that Vps37 proteins dictate the incorporation of UBAP-1, Mvb12A, and Mvb12B leading to the assembly of distinct ESCRT-I complexes, which could be functionally non-redundant. More specifically, our data indicate a partnering preference of Vps37A for UBAP-1 and Vps37B for Mvb12A. This is in line with previous studies on the ESCRT-I stability, which contradicted the stochastic association of ESCRT-I components (Wunderley et al., 2014). We extended these studies showing that concurrent knockdown of all *VPS37* paralogs leads to the nearly complete destabilization of remaining ESCRT-I subunits, resembling effects achieved upon either *TSG101* or *VPS28* silencing. It also explains why we observe similar effects on cell homeostasis, namely induction of inflammatory response and cell growth inhibition, upon knockdown of *TSG101* and concurrent silencing of *VPS37* paralogs and overall similarities in transcriptional responses (Brankatschk et al., 2012; Maminska et al., 2016; Miller et al., 2018). The degree of ESCRT-I destabilization after (co-)silencing of *VPS37* paralogs correlates with the type and magnitude of transcriptional responses; however, the precise mechanism remains to be determined. Destabilization of Vps37 proteins has been well documented upon Tsg101 depletion (Bache et al., 2004; Stefani et al., 2011; Stuchell et al., 2004; Wunderley et al., 2014). Here, we observed mild but distinct transcriptional alterations after either Vps37B or Vps37C depletion that may result from their partly overlapping functions. Both proteins possess the PRR domain and may share similar binding partners. Indeed, using the BioGRID database (Oughtred et al., 2019) we found that both Vps37B and Vps37C have large and partially overlapping interactomes, whilst Vps37A bereft of PRR has only a few interacting proteins. Thus, *VPS37A* as the only paralog encoding the UEV domain might execute functions that cannot be taken over by any other family member. As a consequence, its depletion induces a distinct set of genes than those expressed after knockdown of either *VPS37B* or *VPS37C*. Noteworthy, at least under some conditions, the cell can very well compensate for the loss of a single *VPS37* paralog as illustrated by our RNA-Seq analysis. On the other hand, concurrent silencing of either *VPS37AB* or *VPS37ABC* evokes profound transcriptional alterations that we believe, by similarity to Tsg101 or Vps28 depletion, arise from adverse effects of non-degraded plasma membrane proteins and alterations in protein networks that may contribute to prolonged oncogenic signaling (Maminska et al., 2016). In the context of cancer, ESCRT-I destabilization after concurrent Vps37 depletion could result in more adverse tumor phenotype. This notion is consistent with our transcriptomic analysis of CRC cells depleted of Vps37 proteins, which identified processes related to cell migration, growth and signaling. Downregulation of ESCRT-I components was shown *in vitro* to prolong epidermal growth factor signaling, sensitize cells to low doses of transforming growth factor, as well as promote cell invasion and migration through the process of epithelial to mesenchymal transition (Miller et al., 2018; Yang et al., 2016).

In summary, we established that ESCRT-I subunit destabilization after co-depletion of Vps37 proteins evokes profound cellular stress manifested by a sterile inflammatory response and cell growth arrest. Our findings also revealed potential vulnerabilities of CRC cells with reduced levels of *VPS37A* and *VPS37B* that may be more susceptible to chemotherapeutics and pharmacological modulators of inflammatory response. We also identified candidates with known functions in endocytosis, beyond *VPS37* paralogs, whose expression is changed in CRC and thus warrants further investigation in the context of cancer cell pathophysiology.

## MATERIAL AND METHODS

### Cell culture

Human DLD1 (CCL-221) and RKO (CRL-2577) cell lines were obtained from American Type Culture Collection (ATCC). DLD1 were cultured in Dulbecco’s modified Eagle’s medium (DMEM, Sigma-Aldrich, M2279) supplemented with 10% (v/v) fetal bovine serum (FBS, Sigma-Aldrich, F7524) and 2 mM L-Glutamine (Sigma-Aldrich, G7513). RKO were maintained in Eagle’s minimum essential medium (EMEM, ATCC, 30-2003) supplemented with 10% (v/v) FBS. Both cell lines were passaged using 0.05% Trypsin+EDTA (Sigma-Aldrich, T4049). Cells were cultured in an incubator at 37°C in a humidified atmosphere containing 5% CO_2_. During the study, cells were regularly tested for mycoplasma and the identities of DLD1 and RKO were confirmed by short tandem repeat (STR) profiling performed by the ATCC Cell Authentication Service.

### Cell transfection

Cells were either forward-or reverse-transfected with siRNAs using Lipofectamine RNAiMAX transfection reagent according to the manufacturer’s instructions (Thermo Fisher Scientific, 13778150). The concentration of single siRNA duplex used for transfection was 20 nM. In experiments with simultaneous knockdown of two, three, and four genes, the total concentration of siRNA was 40 nM, 60 nM, and 80 nM, respectively and the proportion of individual siRNAs duplexes was kept equal. The following PreDesigned or Validated Ambion Silencer Select siRNAs (Thermo Fisher Scientific) were used: Negative Control No. 1 (siNC#1, 4390843) and Negative Control No. 2 (siNC#2, 4390846); on-target siVPS37A#1 (s44037), siVPS37A#2 (s44038), siVPS37B#1 (s36177), siVPS37B#2 (s36178), siVPS37C#1 (s30059), siVPS37C#2 (s30060), siRELA (s11916), siCHUK (s3066), siIKBKB (s7263), siCDKN1A#1 (s415), siCDKN1A#2 (s417), siTSG101#1 (s14439), siTSG101#2 (s14440), siVPS28#1 (s27577), siVPS28#2 (s27579), siUBAP1#1 (s27812), siUBAP1#2 (s27813), siMVB12A#1 (s41121), siMVB12A#2 (s41122), siMVB12B#1 (s40157), siMVB12B#2 (s40158). Additionally, two custom-ordered Silencer Select duplexes were used: Negative Control No. 3 (NC3, sense strand 5’->3’ UACGACCGGUCUAUCGUAGtt, antisense strand 5’->3’ CUACGAUAGACCGGUCGUAtt) and Negative Control No. 4 (NC4, sense strand 5’->3’ UUCUCCGAACGUGUCACGUtt, antisense strand 5’->3’ ACGUGACACGUUCGGAGAAtt). The composition of siRNA mixes in experiments with individual and concurrent gene silencing is listed in Table S7.

### Transcriptome analysis by RNA sequencing (RNA-Seq)

Cells were plated in 12-well plate format at the density of 60,000 cells/ml in 1 of medium. After 16-24 h cells were left non-transfected or differentially transfected according to the forward transfection protocol. 72 h later, cells were washed with PBS, and the cell pellet was collected. Sequencing libraries were generated using Ion AmpliSeq Transcriptome Human Gene Expression Panel (ThermoFisher Scientific). Sequencing was performed using Ion Proton instrument with 7 or 8 samples per chip with Ion PI Hi-Q Sequencing 200 Kit (ThermoFisher Scientific). Reads were aligned to the hg19 AmpliSeq Transcriptome ERCC v1 with Torrent Mapping Alignment Program (version 5.0.4, ThermoFisher Scientific). Transcripts were quantified with HTseq-count (version 0.6.0) run with default settings (Anders et al., 2015).

Gene level differential expression analysis was performed using the R package DESeq2 (version 1.18.1; (Love et al., 2014)) for genes with at least 100 counts across conditions and by taking into the account the batch effect and applying the following contrasts (α = 0.05): NT (non-transfected) versus siCTRL#1, NT versus siVPS37A#1, NT versus siVPS37B#1, NT versus siVPS37C#1, NT versus siVPS37AB#1, NT versus siVPS37AC#1, NT versus siVPS37BC#1, NT versus siVPS37ABC#1, siCTRL#1-T versus siVPS37A#1, siCTRL#1-T versus siVPS37B#1, siCTRL#1-T versus siVPS37C#1, siCTRL#1-T versus siVPS37AB#1, siCTRL#1-T versus siVPS37AC#1, siCTRL#1-T versus siVPS37BC#1, siCTRL#1-T versus siVPS37ABC#1. We excluded non-protein coding genes from downstream analysis.

The overlap for different silencing conditions and normalization contrasts was visualized using the VennDiagram package (version 1.6.20). The genes, which overlapped for on-target siRNAs normalized against either NT or siCTRL#1-transfected patterns, were subjected to GO analysis of biological processes and Reactome pathway analysis using clusterProfiler (version 3.6.0; (Yu et al., 2012)) and ReactomePA R-packages (version 3.8; (Yu and He, 2016)) taking advantage of enrichGO and enrichPathway functions, respectively. All enrichment p-values in GO analysis were corrected for multiple testing using the Benjamini-Hochberg method and only genes with adjusted p-value <0.05 were considered significant. The minimal and maximal sizes of gene clusters were set to 10 and 500, respectively. Redundant terms were removed by means of the simplify function with cutoff 0.6. Count data were transformed using the Transcript Per Million (TPM) and scaled across conditions (Z-score). Differentially expressed genes binned in the selected GO processes were used for hierarchical clustering, which was performed on Euclidean distances using Ward’s algorithm. Heatmaps of differentially expressed genes were visualized using ComplexHeatmap (version 1.17.1; (Gu et al., 2016)). All calculations were performed in R version 3.4.4 (https://www.R-project.org).

The code for the present analysis is available on GitHub (https://github.com/kkolmus/VPS37_RNA-Seq). RNA-Seq data have been deposited at Gene Expression Omnibus (GEO) under the accession code: GSE152195.

### Clonogenic assay

Non-transfected cells or cells subjected to reverse transfection with different siRNAs were seeded at the density of 1000 cells per well in a 6-well plate format and cultured for 14 days to form colonies. For staining, colonies were washed with PBS, fixed for 5 min in a 3:1 (v/v) solution of acetic acid:methanol, and incubated for 15 min in 0.2% crystal violet solution in 70% ethanol. The whole procedure was performed at room temperature. Plates with colonies were scanned using the Odyssey Infrared System (LI-COR, Biosciences). Acquired images were analyzed as described before (Guzman et al., 2014). Data are expressed as the percentage staining intensity displayed by non-transfected cells.

### Proliferation assay

1500 cells were left non-transfected or reverse-transfected with different siRNAs in a 96-well plate format and let proliferate for 120 h. BrdU Cell Proliferation ELISA assay (Roche, 11647229001) was used to assess the proliferation of RKO and DLD1 cells according to the manufacturer’s instructions with the following modifications: BrdU reagent was added 4 h prior to cell fixation, 100 µl of substrate solution was added for 5 min followed by addition of 50 µl of 1 M HCl. The colorimetric signal was detected at 450 nm using the Tecan Sunrise Microplate Reader system with the Magellan v. 6.6 software. Data are expressed as the percentage of proliferating, non-transfected cells.

### Western blotting and densitometry analysis

Cells were plated in either 6-or 12-well plate format at the density of 60,000 cells/ml in 1 and 2 ml of medium, respectively. After 16-24 h cells were left non-transfected or differentially transfected according to the forward transfection protocol. 72 h later, cells were lysed in RIPA buffer (1% Triton X-100, 0.5% sodium deoxycholate, 0.1% SDS, 50 mM Tris pH 7.4, 150 mM NaCl, 0.5 mM EDTA) supplemented with protease inhibitor cocktail (6 µg/ml chemostatin, 0.5 µg/ml leupeptin, 10 µg/ml antipain, 2 µg/ml aprotinin, 0.7 µg/ml pepstatin A, 10 µg/ml 4-amidinophenylmethanesulfonyl fluoride hydrochloride; Sigma-Aldrich) and phosphatase inhibitor cocktails 2 and 3 (P5726 and P0044, Sigma-Aldrich). Protein concentration was determined using the BCA Protein Assay Kit (Thermo Fisher Scientific, 23225). 25-30 µg of total protein per sample was resolved on 12 or 15% SDS-PAGE, transferred onto a nitrocellulose membrane (Amersham Hybond, GE Healthcare Life Science, 10600002), blocked with 5% milk in PBS with 1% Tween, probed first with specific primary and then secondary antibodies, and imaged using the detection solution (BioRad, 170-5061) and ChemiDoc imaging system (Bio-Rad). All primary antibodies are listed in Table S8. Secondary horseradish peroxidase-conjugated anti-mouse (315-005-008), anti-rabbit (111-035-144) and anti-goat (305-035-046) antibodies were from Jackson ImmunoResearch and were used at working dilution 1:10,000. Densitometry of protein bands was carried out using ImageJ software (Schneider et al., 2012). p65 was used as the loading control for quantification of phosphorylated p65. Vinculin was used as a loading control in all other experiments. Results are presented as fold change compared to non-transfected cells.

### Quantitative Real-Time Polymerase Chain Reaction (qRT-PCR)

Cells were seeded in a 12-well plate format at the density of 60,000 cells/ml in 1 ml of medium. After 16-24 h cells were left non-transfected or differentially transfected according to the forward transfection protocol. 72 h later, total RNA was isolated with High Pure Isolation Kit (Roche, 11828665001). 500 ng of total RNA was subjected for cDNA synthesis. M-MLV, random nonamers and oligo(dT)_23_ (Sigma-Aldrich, M1302, R7647, and O4387, respectively) were used for cDNA synthesis according to the manufacturer’s instructions. Expression of genes of interest was measured using primers designed with the NCBI Primer designing tool and custom-synthesized by Sigma-Aldrich. We list primers used in the present study (Table S9). Real-Time cDNA amplification was performed with the Kapa Sybr Fast qPCR Kit (KapaBiosystems, KK4618). Fluorescence was monitored using the 7900HT Fast Real-Time PCR thermocycler (Applied Biosystems). Expression of each gene was normalized to either expression of the *ACTB* (β-actin) or *GAPDH* (glyceraldehyde 3-phosphate dehydrogenase) reference genes. Results are presented as fold change compared to non-transfected cells. For clarity, the Y-axis is interrupted in some cases.

### Analysis of expression levels of VPS37 paralogs in healthy and colorectal cancer (CRC) samples using qRT-PCR

Samples of the normal colon (n=24) and adenocarcinoma (n=26) had been collected for the purpose of previous studies (Mikula et al., 2011; Skrzypczak et al., 2010). In order to determine the abundance of *VPS37A* and *VPS37B* transcripts, qRT-PCR was performed as described before (Mikula et al., 2011; Skrzypczak et al., 2010). The sequences of primers for *VPS37A* and *VPS37B* are listed in Table S9.

### Flow cytometry analysis

Cells were seeded in a 6-well plate format at the density of 60,000 cells/ml in 2 ml of medium. After 16-24 h cells were left non-transfected or differentially transfected according to the forward transfection protocol. 96 h post-transfection cells were briefly washed with PBS, harvested with trypsin+EDTA, washed twice with PBS, and fixed for 24 h in ice-cold 70% ethanol. Washed cells were then incubated first with extraction buffer (4 mM citric acid in 0.2 M Na_2_HPO_4_) for 5 min at room temperature, and next with staining solution (3.8 mM sodium citrate, 50 µg/ml propidium iodide (PI) and 0.5 mg/ml RNase A) for 30 min at room temperature. Analysis of cells was performed on the BD LSRFortessa flow cytometer (Bekton Dickinson). A total of 10,000 cells from single cell gate were counted for each transfection condition. Flow cytometry data were plotted and analyzed by FlowJo (Bekton Dickinson) and ModFit LT (Verity Software House) software. Data is presented as percentage of analyzed cells in the given cell cycle phase.

### Immunohistochemistry (IHC) and analyses of normal and CRC samples

The study protocol for analysis of protein levels of Vps37A and Vps37B in human normal colon and CRC samples was approved by the Bioethics Committee of the Maria Skłodowska-Curie National Research Institute of Oncology in Warsaw (decision no. 40/2017). Informed consent was obtained from all subjects. The experiment conformed to the principles set out in the WMA Declaration of Helsinki and the Department of Health and Human Services Belmont Report. High-density tissue microarrays were constructed from formalin-fixed, paraffin-embedded diagnostic samples of 100 pairs of treatment-naïve CRC tissues and matched normal colon samples from the collection of the Maria Skłodowska-Curie National Research Institute of Oncology in Warsaw. IHC was performed using automated immunohistochemical stainer (Dako Denmark A/S) and specific anti-VPS37A and anti-VPS37B antibodies listed in Table S8. The EnVision Detection System (Agilent) was used for detection. Samples were reviewed for the abundance of Vps37 proteins in normal and neoplastic tissue by two pathologists who were blinded to the outcome. A semi-quantitative method was applied for IHC evaluation, involving a scoring system based on the staining intensity: 0 – no staining, 1+ – weak, 2+ – intermediate, 3+ strong staining. Staining homogeneity was above 90%.

### TCGA data analysis

Clinicopathological and transcriptional profiles from the two TCGA cohorts: rectum adenocarcinoma (READ) and colon adenocarcinoma (COAD) were retrieved using the TCGAbiolinks package (Colaprico et al., 2016). READ and COAD datasets were analyzed together as a previous study showed a major overlap in their expression patterns (Weinstein et al., 2013). The present analysis encompassed only matched normal-tumor tissue samples for which clinicopathological data were available. 31 patients fulfilled the latter criterion. 19 of these patients were assigned to the early stage group encompassing stages I and II, and 12 were assigned to the advanced stage group encompassing stages III and IV. Matching TCGA sequencing data acquired using the Illumina HiSeq platform were used. Differential gene expression was performed with the default settings on count data taking into account correction of batch effect relying on the inter-plate variation. Only transcripts whose gene counts exceeded 50 were selected for downstream analysis. Volcano plots of differentially expressed genes were prepared with ggplot2 (CRAN available package). Heatmaps of differentially expressed genes were visualized using ComplexHeatmap (version 1.17.1; (Gu et al., 2016)). The overlap between differentially expressed genes was visualized using the VennDigram package (CRAN available package). All calculations were performed in R version 3.6.1 (https://www.R-project.org). The code for the present analysis is available on GitHub (https://github.com/kkolmus/TCGA_transcriptomic_proj).

### Statistical analysis

Data are shown as mean ± standard deviation of at least three independent biological experiments. For qRT-PCR and BrdU, samples were assayed in technical triplicates. Statistical analysis was performed using the GraphPad Prism 6 software. Experiments with normal distribution were analyzed using either Student’s t-test or one-way ANOVA followed by Bonferroni’s multiple comparison test. In the case of non-normal distribution Mann-Whitney U-test was applied. Analysis of categorical data was performed using Fisher’s exact test. The significance of mean comparison is annotated as follows: ns, non-significant (*P*≥0.05), \**P*<0.05, \*\**P*<0.01, \*\*\**P*<0.001, \*\*\*\**P*<0.0001. Results were considered significant when *P*<0.05. No statistical methods were used to predetermine sample size.

## Acknowledgments

We thank A. Zeigerer (Helmholtz Zentrum Munchen), D. Zdzalik-Bielecka, J. Cendrowski, A. Poswiata, and M. Kaczmarek for critical reading of the manuscript. We also thank D. Zdzalik-Bielecka (International Institute of Molecular and Cell Biology) for help with pilot FACS experiments. We are grateful to A. Paziewska and A. Dabrowska (Maria Sklodowska-Curie National Research Institute of Oncology) for their technical support in RNA-Seq analysis.

## Author contributions

The research was conceived by M. Miaczynska, K.K. and M. Mikula. Funding was acquired by M. Miaczynska. Experiments were designed and performed mostly by K.K. P.E., B.S. and E.S., with input from M. Miaczynska, E.S. and crucial help from M. Mikula and K.G. (RNA-Seq), E.D-W, A.S-C. and M.P-S. (immunohistochemistry), and MB-O. and K.P. (flow cytometry). The manuscript was written by K.K. and M. Miaczynska. Figures were assembled by K.K. with help of A.S-C. K.K. and M. Miaczynska supervised the work. All authors approved the manuscript.

## Data and materials availability

The RNA-Seq datasets have been deposited to GEO under the accession number: GSE152195 (https://www.ncbi.nlm.nih.gov/geo/query/acc.cgi?acc=GSE152195).

---

**Fig. S1.**
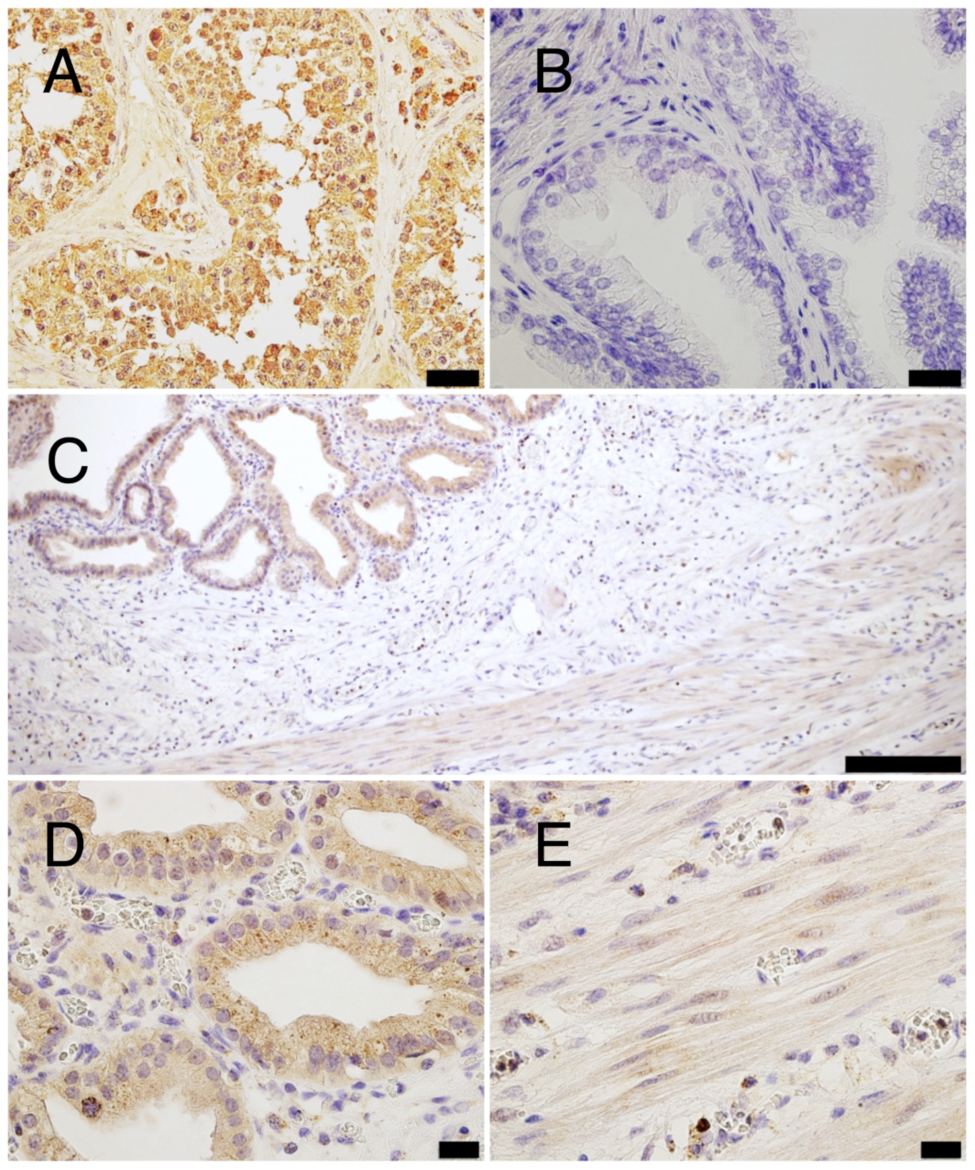
Immunohistochemical evaluation of the specificity of antibodies against Vps37A and Vps37B proteins. Vps37A: (A) positive control - strong granular cytoplasmic staining in the glandular epithelium of testis (scale bar 20 µm), (B) lack of staining in the glandular epithelium of prostate (scale bar 10 µm). **Vps37B:** (C) positive control - strong granular cytoplasmic staining in the mucosa of the gallbladder (scale bar 100 µm) and (D) at higher magnification (scale bar 10 µm), (E) weak staining in the muscle (scale bar 10 µm).

**Fig. S2.**
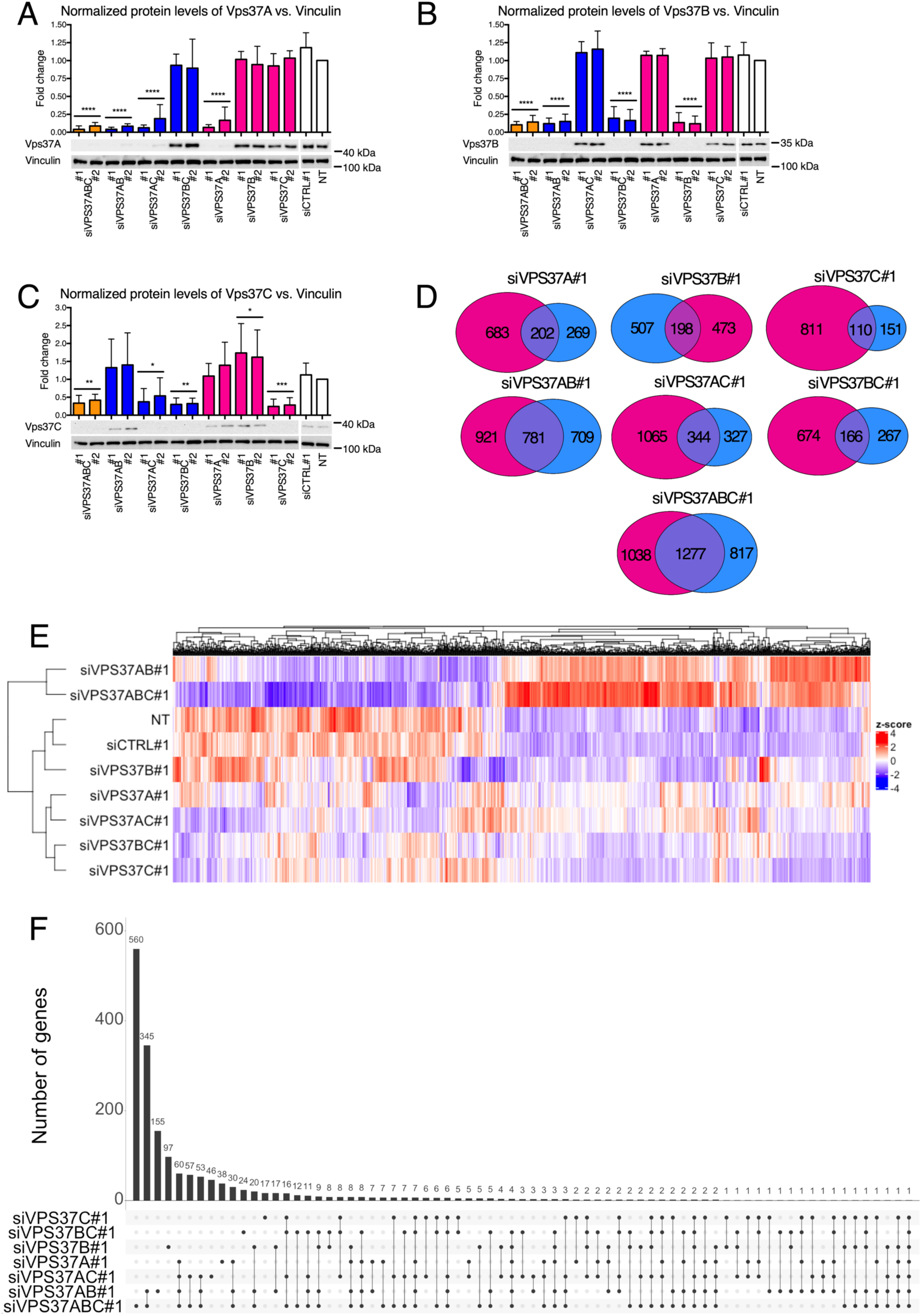
Transcriptional alteration after individual or combined knockdown of *VPS37* paralogs in DLD1 cells. (A-C) Western blotting analysis of selectivity of siRNAs for the *VPS37* paralogs: (A) *VPS37A*, (B) *VPS37B* and (C) *VPS37C*. Lysates were collected 72 h after transfection with siRNA targeting individual or combinations of *VPS37* paralogs. Lysates from non-transfected (NT) cells and transfected with non-targeting siRNA (siCTRL#1) were used to assess the basal level of intracellular signaling. Vinculin was used as a loading control. Representative blots from n=5 independent experiments are shown along with densitometry analysis. Data are mean ± standard deviation expressed as the fold change of protein level in NT cells, which was set as 1. Statistical significance was assessed against grouped NT and siCTRL#1 conditions using one-way ANOVA test followed by Bonferroni’s correction; \**P*<0.05, \*\**P*<0.01, \*\*\**P*<0.001, \*\*\*\**P*<0.0001. (D) Venn diagrams of differentially expressed genes (≥ 1.50-fold or ≤ 0.667-fold; adjusted P<0.05) after single, double or triple *VPS37* paralog silencing when normalized to non-transfected (NT, pink circles) cells and cells transfected with non-targeting siRNA (siCTRL#1, blue circles). Differentially expressed genes were identified using DESeq2. (E) Heatmap visualizing clustering of expression patterns for differentially expressed genes in DLD1 cells across different transfection conditions. (F) Clustered heatmap showing pairwise intersections for the common pools of differentially expressed genes (≥ 1.50-fold or ≤ 0.667-fold; adjusted *P*<0.05) after single, double or triple *VPS37* paralog silencing when normalized to non-transfected cells and cells transfected with non-targeting siRNA for the investigated transfection conditions. RNA-Seq data analysis was performed using n=3 independent experiments.

**Fig. S3.**
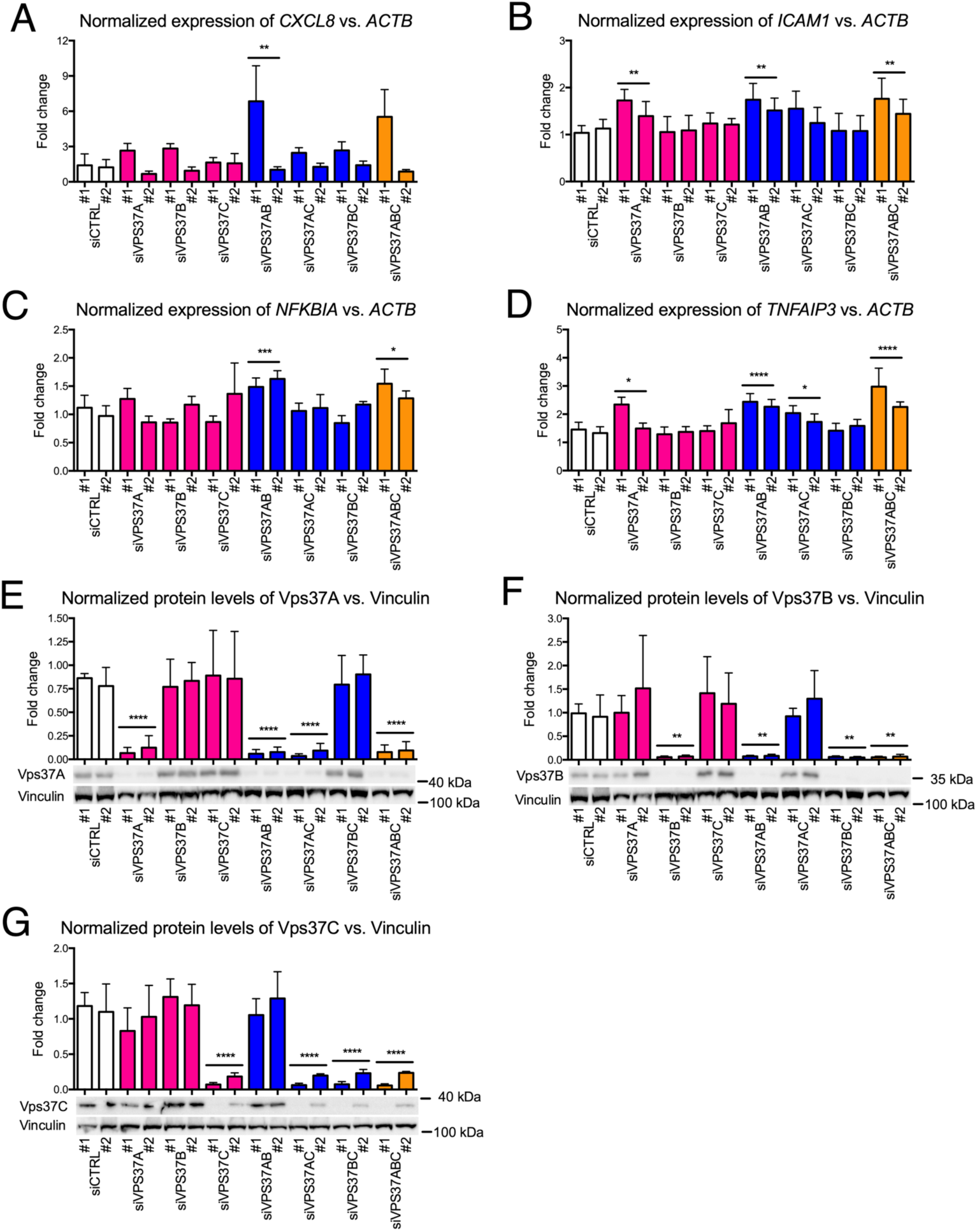
Induction of inflammatory gene expression upon concurrent silencing of *VPS37* paralogs. (A-D) qRT-PCR analysis of selected genes in the inflammatory response cluster: (A) *CXCL8*, (B) *ICAM1*, (C) *NFKBIA* and (D) *TNFAIP3* in DLD1 cells. Expression of genes was measured 72 h after forward transfection with siRNA targeting of *VPS37* paralogs individually or in combinations. Non-transfected (NT) cells and transfected with non-targeting siRNA (siCTRL) were used to assess the basal expression level of the investigated genes. *ACTB* (encoding β-Actin) was used as a reference gene. Data in panels (A-D) are mean ± standard deviation of n=5 independent experiments. Data are expressed as the fold change of mRNA levels in NT cells, which was set as 1. (E-G) Western blotting analysis of selectivity of siRNAs for the *VPS37* paralogs. Lysates were collected 72 h after forward transfection with siRNA targeting individual or combinations of *VPS37* paralogs. Lysates from NT cells and transfected with non-targeting siRNA (siCTRL#1 and siCTRL#2) were used to assess the basal level of intracellular signaling. Vinculin was used as a loading control. Representative blots from n=3 independent experiments are shown along with densitometry analysis. Data are mean ± standard deviation expressed as the fold change of protein level in NT cells, which was set as 1. Statistical significance in all panels was assessed against grouped siCTRL conditions using one-way ANOVA test followed by Bonferroni’s correction; \**P*<0.05, \*\**P*<0.01, \*\*\**P*<0.001, \*\*\*\**P*<0.0001.

**Fig. S4.**
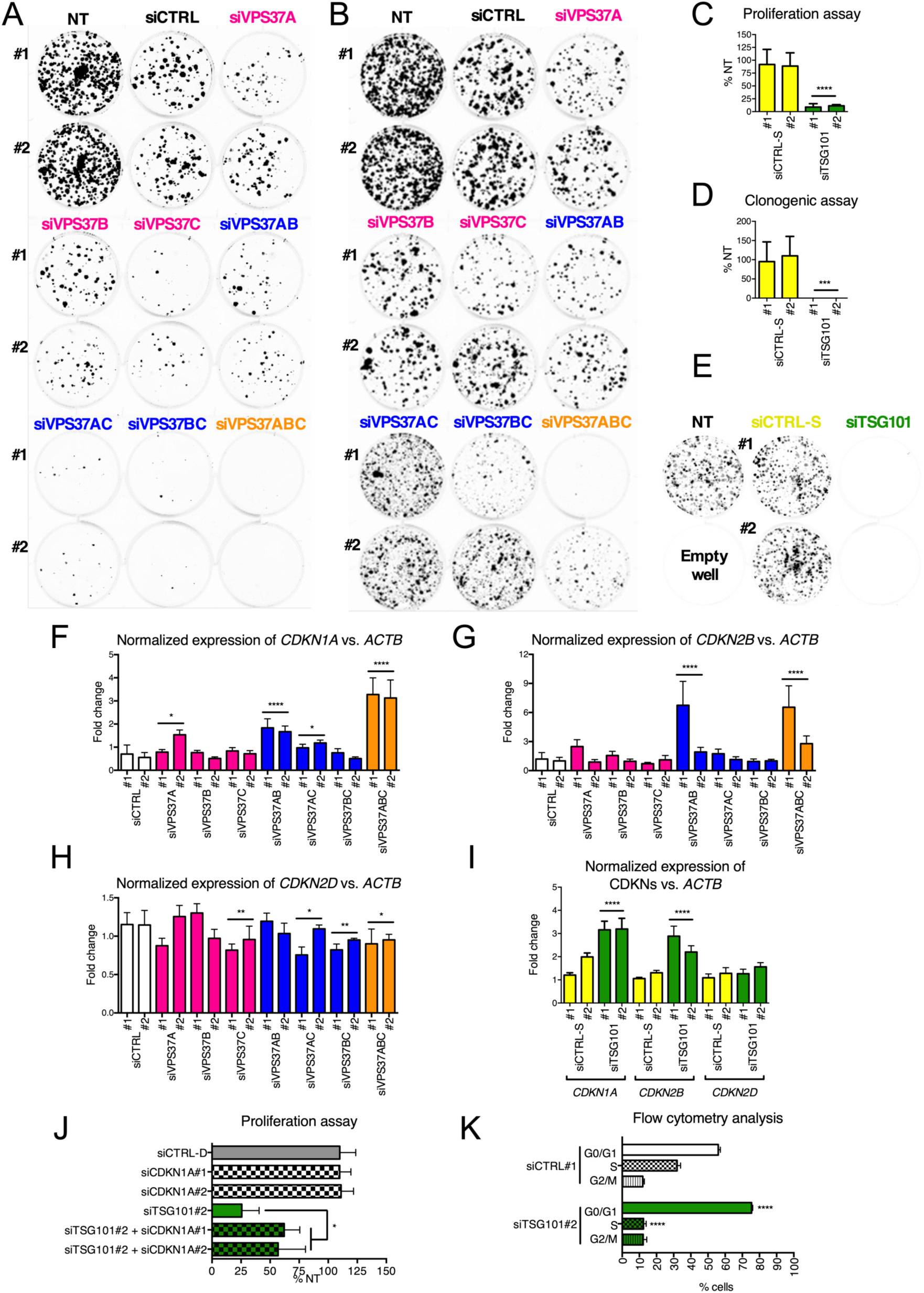
CRC cell growth inhibition upon knockdown of all *VPS37* paralogs or *TSG101*. (A, B) Representative images of clone formation assessed 14 days after differential knockdown of *VPS37* paralogs in (A) DLD1 and (B) RKO cells. (C) Cell proliferation of RKO cells assessed 120 h after *TSG101* knockdown using the BrdU proliferation assay. (D) Clonogenic growth of RKO cells assessed 14 days after *TSG101* knockdown. (E) Representative images of clone formation assessed 14 days after knockdown of *TSG101* in RKO cells. (F-H) qRT-PCR analysis of expression of genes encoding CDKNs: (F) *CDKN1A*, (G) *CDKN2B* and (H) *CDKN2D* in DLD1 cells. Expression of *CDKNs* was measured 72 h after individual and concurrent depletion of Vps37 proteins in n=5 independent experiments. (I) qRT-PCR analysis of expression of genes encoding CDKNs: *CDKN1A, CDKN2B* and *CDKN2D* in RKO cells. Expression of *CDKNs* was measured 72 h after Tsg101 depletion. Non-transfected (NT) cells and transfected with non-targeting siRNA (siCTRL) were used to assess the basal level of the investigated genes. *ACTB* (encoding β-Actin) was used as a reference gene. (J) Cell proliferation of RKO cells was assessed 120 h after concurrent silencing of *TSG101* and *CDKN1A* using the proliferation assay. (K) Analysis of cell cycle was performed upon Tsg101 depletion. Cells were forward transfected for 96 h, stained with PI and evaluated with a flow cytometer. Unless stated otherwise data presented in all panels are mean of n=3 independent experiments ± standard deviation analyzed with one-way ANOVA with Bonferroni’s correction (F-H, J) or Student’s t-test (C, D, I, K). Statistical significance for grouped siCTRL conditions (A-J) and for siCTRL#1 in panel K, * *P*<0.05; ** *P*<0.01, *** *P*<0.001, **** *P*<0.0001.

**Fig. S5.**
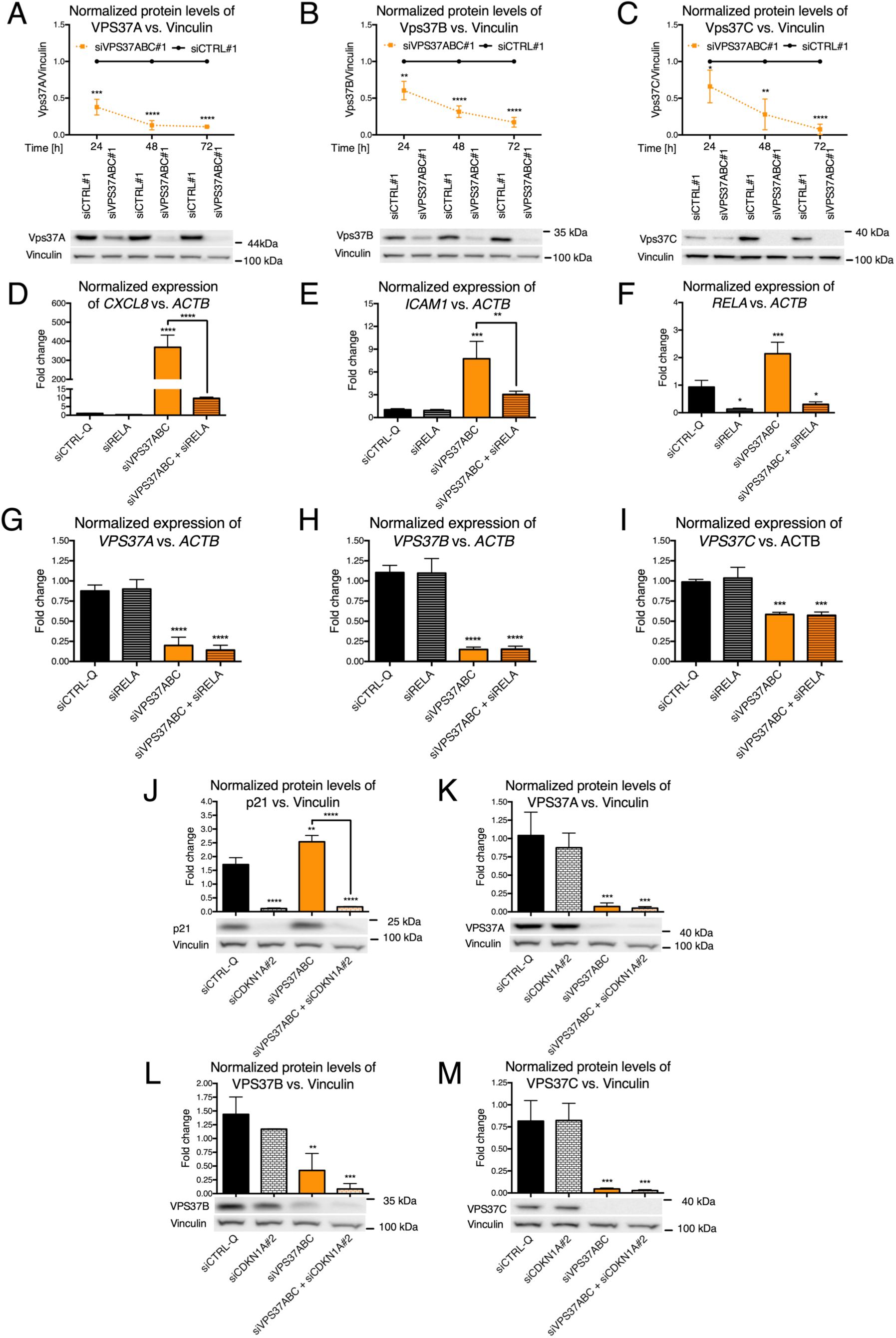
Control experiments corroborating independent activation of NF-κB inflammatory response and p21-mediated inhibition of cell growth. (A-C) Western blot analyses of (A) Vps37A, (B) Vps37B, and (C) Vps37C levels were performed 24, 48 and 72 h after transfection with non-targeting or on-target siRNA for *VPS37ABC*. (D-K) qRT-PCR analyses of (D) *CXCL8*, (E) *ICAM1*, (F) *RELA*, (G) *VPS37A*, (H) *VPS37B, and* (I) *VPS37C* and (I) were performed 72 h after transfection with siRNA targeting the p65 component of NF-κB heterodimers alone or in combination with the three *VPS37* paralogs. (J-M) Western blotting analyses of (J) p21, (K) Vps37A, (L) Vps37B and (M) Vps37C abundance were performed 72 h after transfection with siRNA targeting *CDKN1A* alone or in combination with the three VPS37 paralogs. (D-K) *ACTB* (encoding β-Actin) was used as a reference gene in qRT-PCR analysis. (A-C, J-M) Vinculin was used as a loading control for Western blotting. Representative blots are shown along with densitometry analysis. Data in all panels are mean ± standard deviation of n=3 independent experiments expressed as the fold change of either mRNA or protein abundance. In panels A-C protein abundance in siCTRL#1 transfected cells was set to 1, whilst in panels D-I mRNA and protein abundance in non-transfected (NT) cells was set as 1. Statistical significance was assessed using unpaired Student’s t-test (A-C) or one-way ANOVA test followed by Bonferroni’s correction (D-M). Statistical significance against siCTRL#1 at matching time point for transfection with three different siRNA (A-C) and siCTRL-Q for four different siRNA (D-M), \**P*<0.05, \*\**P*<0.01, \*\*\**P*<0.001, \*\*\*\**P*<0.0001.

**Fig. S6.**
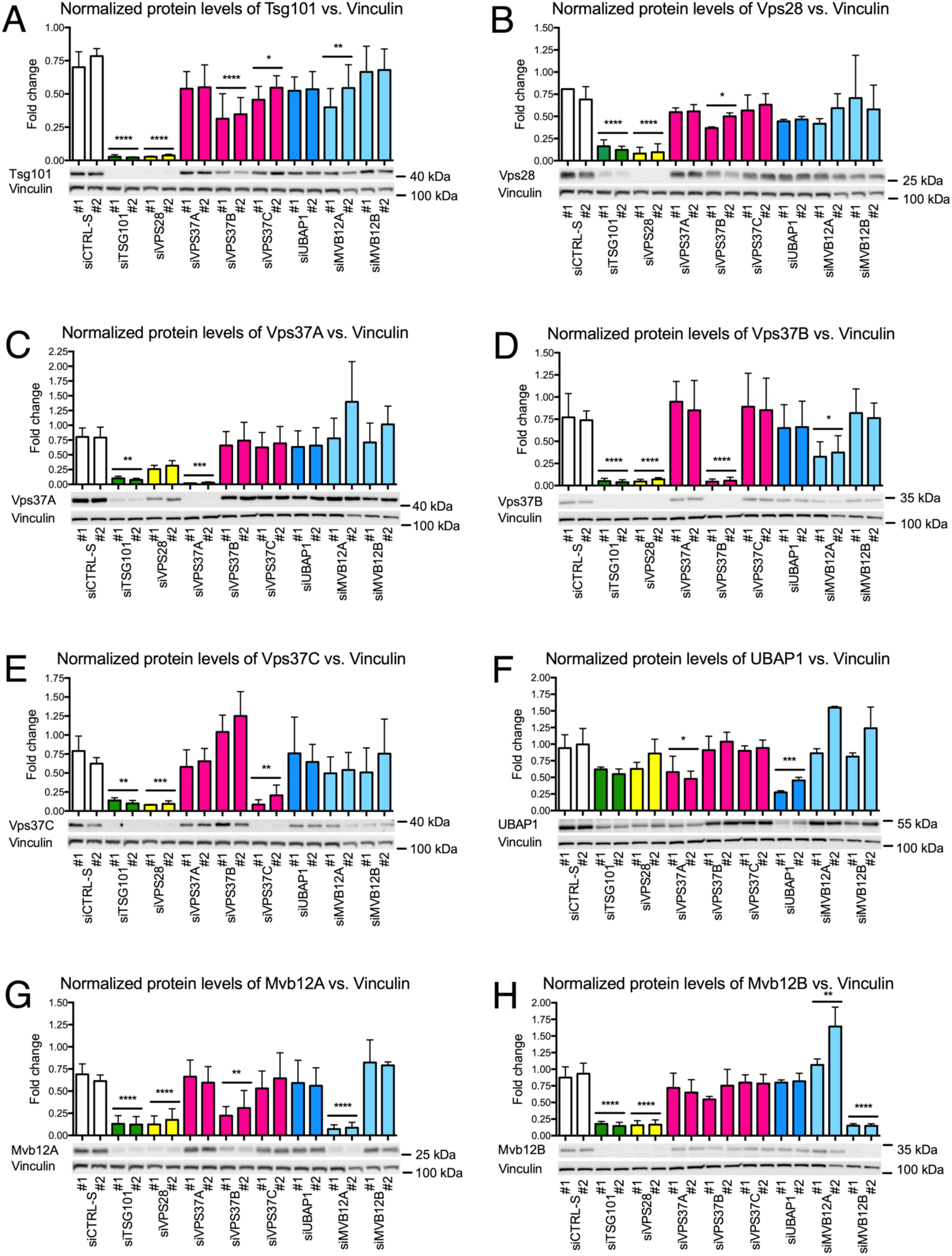
Destabilization of ESCRT-I subunits upon depletion of Tsg101 and Vps28. Western blotting analysis of ESCRT-I subunits: (A) Vps37A, (B) Vps37B, (C) Vps37C, (D) Tsg101, (E) Vps28, (F) UBAP-1, (G) Mvb12A and (H) Mvb12B. Lysates of RKO cells were collected 72 h after transfection with siRNA targeting individual ESCRT-I components. Lysates from non-transfected cells and transfected with non-targeting siRNA were used to assess the basal level of ESCRT-I subunits. Vinculin was used as a loading control. Representative blots are shown along with densitometry analysis. Data are mean ± standard deviation of n=3 independent experiments expressed as the fold change of protein level in non-transfected cells, which was set as 1. Statistical significance was assessed using ANOVA test followed by Bonferroni’s correction. Statistical significance against siCTRL-S conditions, \**P*<0.05, \*\**P*<0.01, \*\*\**P*<0.001, \*\*\*\**P*<0.0001.

**Table S1.**
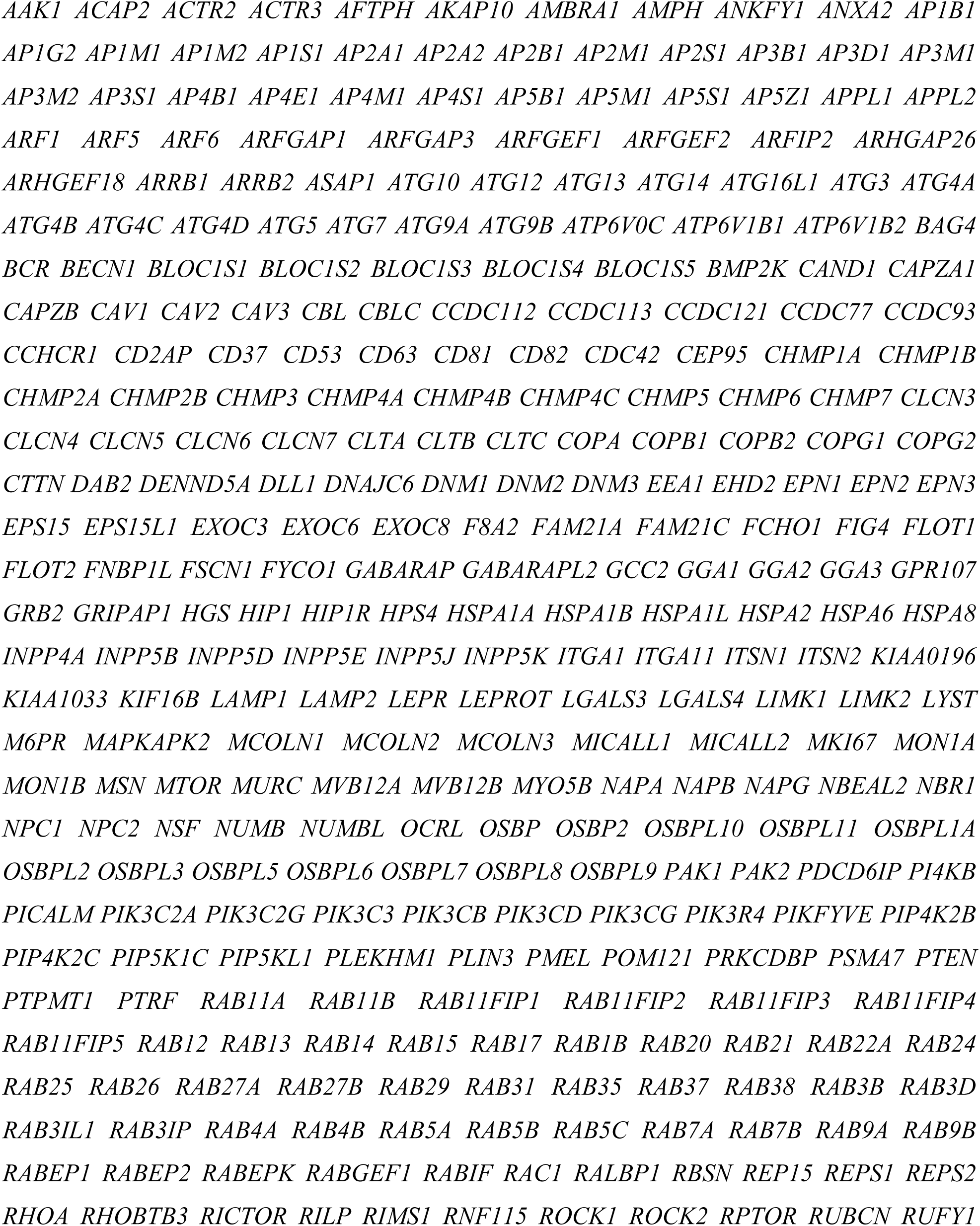
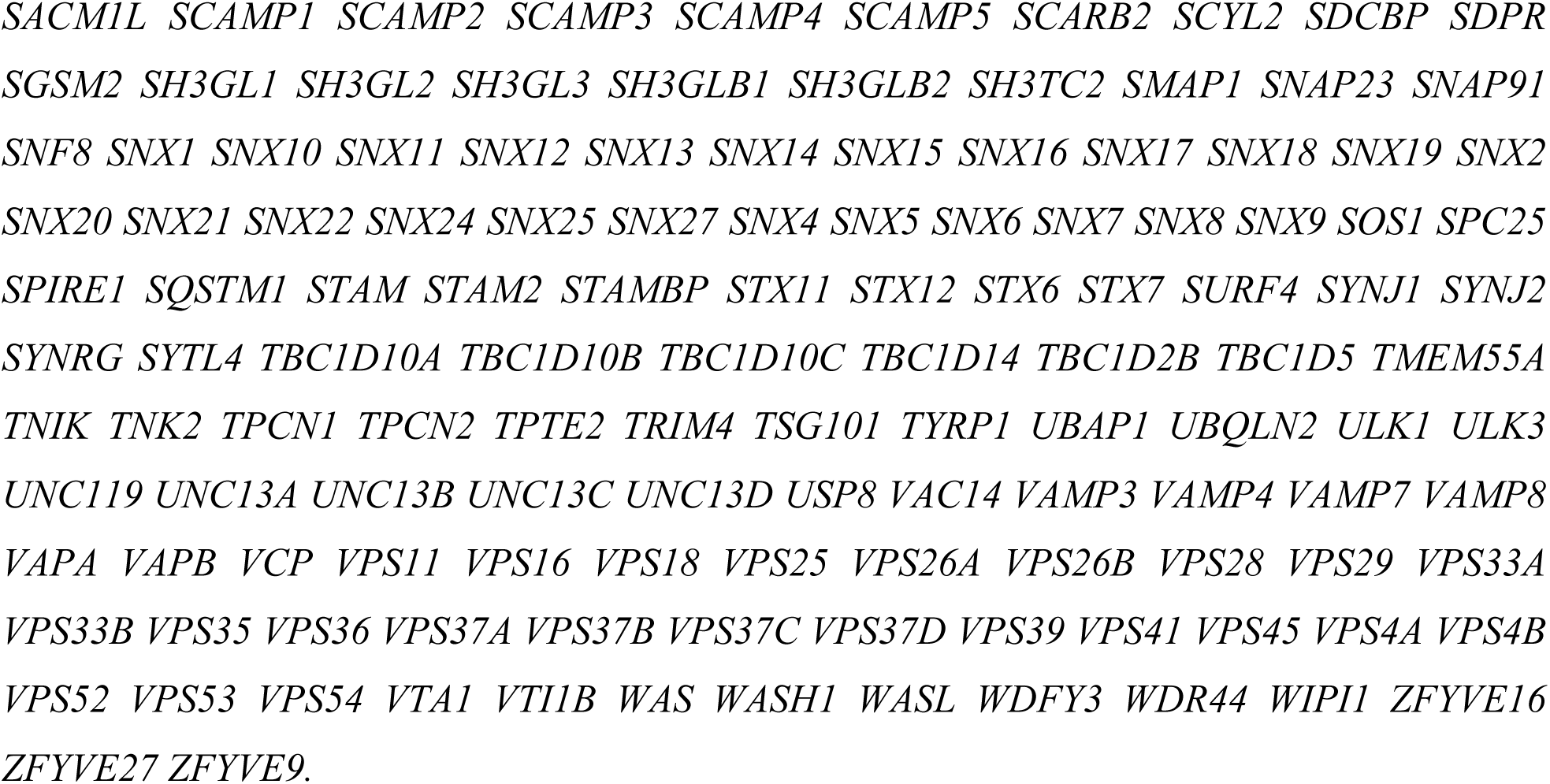
List of genes used for TCGA data mining.

**Table S2.**
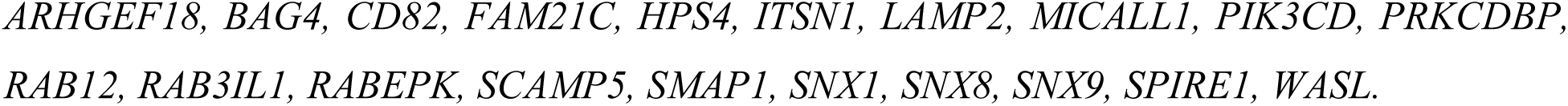
List of differentially expressed genes in the early stages of CRC.

**Table S3.**
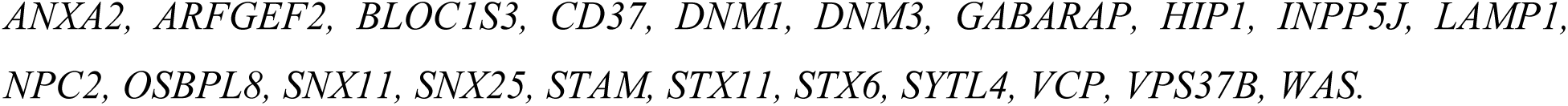
List of differentially expressed genes in the advanced stages of CRC.

**Table S4.** Expression of genes encoding ESCRT-I subunits in COAD and READ cohorts of TCGA patients at the early (stage I and II) and advanced stages (stage III and IV) of CRC. Table shows fold change ratio between the matched healthy and cancer patient tissue in the logarithmic scale (Log2FC); with associated false discovery rate (FDR) upon differential expression analysis using TCGAbiolinks.

| Gene | Early stages |  | Advanced stages |  |
| --- | --- | --- | --- | --- |
|  | Log2FC | FDR | Log2FC | FDR |
| *VPS37A* | -0.294 | 0.09 | -0.308 | 0.23 |
| *VPS37B* | -0.266 | 0.15 | -0.686 | 0.02 |
| *VPS37C* | -0.070 | 0.64 | -0.102 | 0.71 |
| *TSG101* | -0.006 | 0.97 | -0.136 | 0.66 |
| *VPS28* | -0.439 | 0.01 | 0.198 | 0.52 |

**Table S5.** Association of Vps37A and Vps37B protein levels clinicopathological features of n=100 CRC patients. Abbreviations: pT - pathological tumors status, pN – pathological nodes status, G – disease grade.

| pT status | Vps37A staining intensity | Vps37B staining intensity | Number of patients |
| --- | --- | --- | --- |
| 1 | 2 | 2 | 2 |
| 2 | 2 | 1 | 4 |
| 2 | 2 | 2 | 13 |
| 3 | 2 | 1 | 22 |
| 3 | 2 | 2 | 43 |
| 3a | 2 | 2 | 1 |
| 3b | 2 | 1 | 1 |
| 3b | 2 | 2 | 1 |
| 3c | 2 | 1 | 1 |
| 3c | 2 | 2 | 1 |
| 4a | 2 | 1 | 1 |
| 4a | 2 | 2 | 4 |
| 4b | 2 | 1 | 1 |
| 4b | 2 | 2 | 5 |

| pN status | Vps37A staining intensity | Vps37B staining intensity | Number of patients |
| --- | --- | --- | --- |
| 0 | 2 | 1 | 15 |
| 0 | 2 | 2 | 48 |
| 1a | 2 | 1 | 5 |
| 1a | 2 | 2 | 12 |
| 1b | 2 | 2 | 7 |
| 1c | 2 | 1 | 2 |
| 1c | 2 | 2 | 1 |
| 2a | 2 | 1 | 5 |
| 2a | 2 | 2 | 2 |
| 2b | 2 | 1 | 3 |

| G | Vps37A staining intensity | Vps37B staining intensity | Number of patients |
| --- | --- | --- | --- |
| 2 | 2 | 1 | 21 |
| 2 | 2 | 2 | 51 |
| 3 | 2 | 1 | 9 |
| 3 | 2 | 2 | 19 |

**Table S6.** List of differentially expressed genes in COAD and READ cohorts of TCGA patients at the early (stage I and II) and advanced stages (stage III and IV) of CRC. Table is available as a supplemental file for this manuscript. Table shows fold change ratio between the matched healthy and cancer patient tissue in the logarithmic scale (Log2FC); with associated false discovery rate (FDR) upon differential expression analysis using TCGAbiolinks. Genes with increased and decreased expression are those with FDR < 0.05 and log2FC ≥ 0.6 and ≤ −0.6, respectively. Empty records signify no significant changes for the given gene.

**Table S7.**
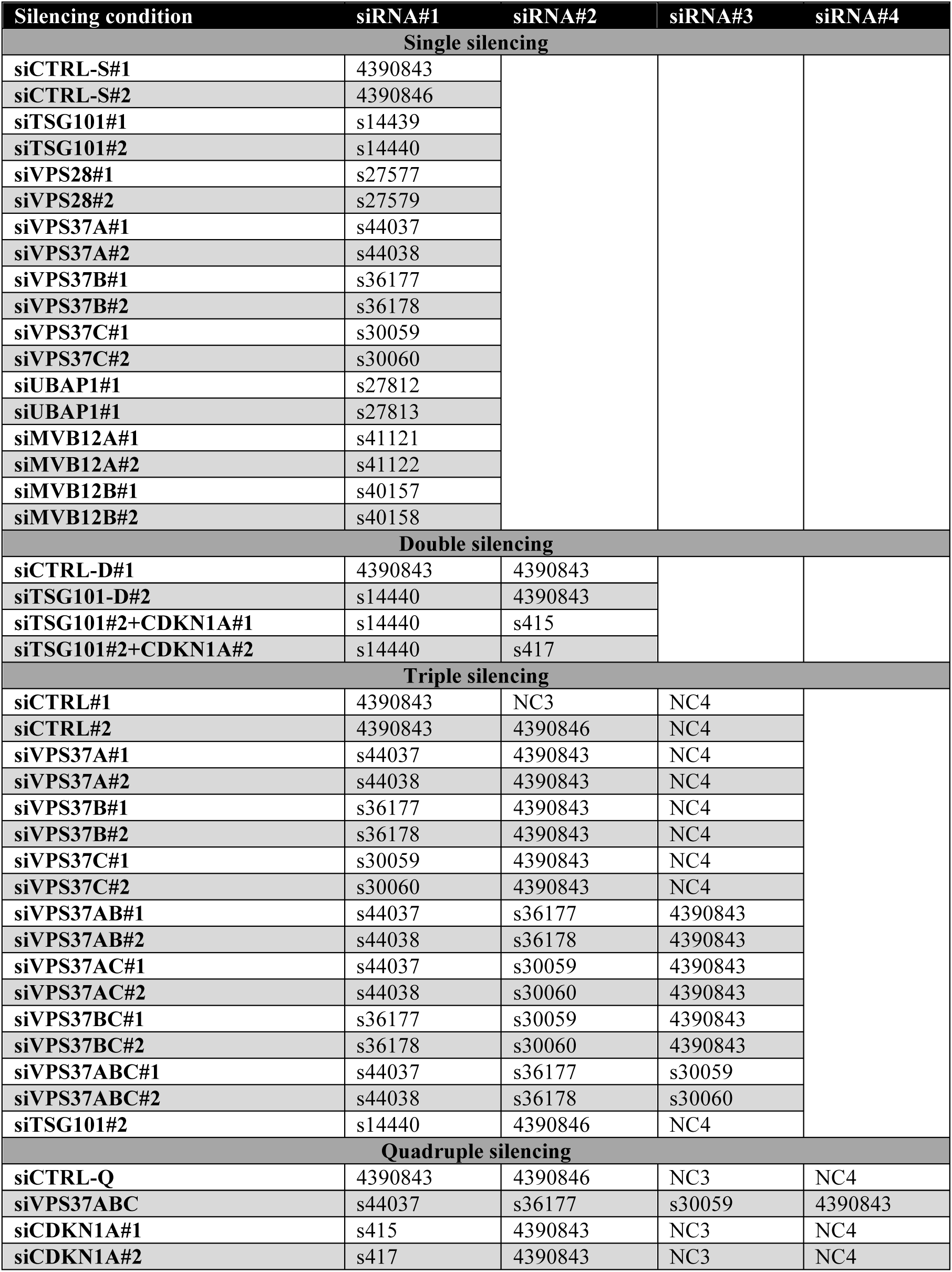
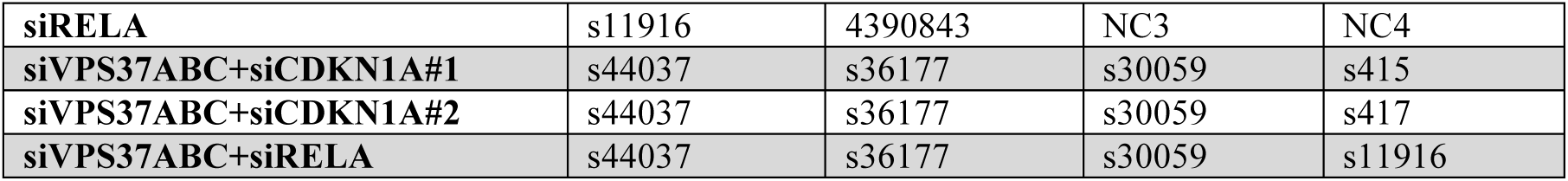
List of siRNA combinations used in the present study.

| Silencing condition | siRNA#1 | siRNA#2 | siRNA#3 | siRNA#4 |
| --- | --- | --- | --- | --- |
| Single silencing |  |  |  |  |
| siCTRL-S#1 | 4390843 |  |  |  |
| siCTRL-S#2 | 4390846 |  |  |  |
| siTSG101#1 | s14439 |  |  |  |
| siTSG101#2 | s14440 |  |  |  |
| siVPS28#1 | s27577 |  |  |  |
| siVPS28#2 | s27579 |  |  |  |
| siVPS37A#1 | s44037 |  |  |  |
| siVPS37A#2 | s44038 |  |  |  |
| siVPS37B#1 | s36177 |  |  |  |
| siVPS37B#2 | s36178 |  |  |  |
| siVPS37C#1 | s30059 |  |  |  |
| siVPS37C#2 | s30060 |  |  |  |
| siUBAP1#1 | s27812 |  |  |  |
| siUBAP1#1 | s27813 |  |  |  |
| siMVB12A#1 | s41121 |  |  |  |
| siMVB12A#2 | s41122 |  |  |  |
| siMVB12B#1 | s40157 |  |  |  |
| siMVB12B#2 | s40158 |  |  |  |
| Double silencing |  |  |  |  |
| siCTRL-D#1 | 4390843 | 4390843 |  |  |
| siTSG101-D#2 | s14440 | 4390843 |  |  |
| siTSG101#2+CDKN1A#1 | s14440 | s415 |  |  |
| siTSG101#2+CDKN1A#2 | s14440 | s417 |  |  |
| Triple silencing |  |  |  |  |
| siCTRL#1 | 4390843 | NC3 | NC4 |  |
| siCTRL#2 | 4390843 | 4390846 | NC4 |  |
| siVPS37A#1 | s44037 | 4390843 | NC4 |  |
| siVPS37A#2 | s44038 | 4390843 | NC4 |  |
| siVPS37B#1 | s36177 | 4390843 | NC4 |  |
| siVPS37B#2 | s36178 | 4390843 | NC4 |  |
| siVPS37C#1 | s30059 | 4390843 | NC4 |  |
| siVPS37C#2 | s30060 | 4390843 | NC4 |  |
| siVPS37AB#1 | s44037 | s36177 | 4390843 |  |
| siVPS37AB#2 | s44038 | s36178 | 4390843 |  |
| siVPS37AC#1 | s44037 | s30059 | 4390843 |  |
| siVPS37AC#2 | s44038 | s30060 | 4390843 |  |
| siVPS37BC#1 | s36177 | s30059 | 4390843 |  |
| siVPS37BC#2 | s36178 | s30060 | 4390843 |  |
| siVPS37ABC#1 | s44037 | s36177 | s30059 |  |
| siVPS37ABC#2 | s44038 | s36178 | s30060 |  |
| siTSG101#2 | s14440 | 4390846 | NC4 |  |
| Quadruple silencing |  |  |  |  |
| siCTRL-Q | 4390843 | 4390846 | NC3 | NC4 |
| siVPS37ABC | s44037 | s36177 | s30059 | 4390843 |
| siCDKN1A#1 | s415 | 4390843 | NC3 | NC4 |
| siCDKN1A#2 | s417 | 4390843 | NC3 | NC4 |
| **siRELA** | s11916 | 4390843 | NC3 | NC4 |
| **siVPS37ABC+siCDKN1A#1** | s44037 | s36177 | s30059 | s415 |
| **siVPS37ABC+siCDKN1A#2** | s44037 | s36177 | s30059 | s417 |
| **siVPS37ABC+siRELA** | s44037 | s36177 | s30059 | s11916 |

**Table S8.** List of antibodies used in the present study. Abbreviations: WB – Western blot, IHC – Immunohistochemistry.

| **Antibody** | **Species** | **Technique** | **Dilution** | **Company** | **Catalog number** |
| --- | --- | --- | --- | --- | --- |
| **VPS37A** | Rabbit | WB, IHC | WB 1:1000<br>IHC 1:100 | Proteintech | 11870-1-AP |
| **VPS37B** | Rabbit | WB, IHC | WB 1:1000<br>IHC 1:100 | Sigma | HPA-038218 |
| **VPS37C** | Goat | WB | WB 1:500 | Abcam | Ab40851 |
| **TSG101** | Rabbit | WB | WB 1:1000 | Abcam | Ab133586 |
| **VPS28** | Rabbit | WB | WB 1:1000 | Abcam | Ab167172 |
| **UBAP1** | Rabbit | WB | WB 1:1000 | Proteintech | 12385-1-AP |
| **MVB12A** | Rabbit | WB | WB 1:1000 | Atlas Antibodies | HPA041885 |
| **MVB12B** | Rabbit | WB | WB 1:1000 | Atlas Antibodies | HPA043683 |
| **Phospho-p65** | Rabbit | WB | WB 1:1000 | Cell Signaling Technology | 3033 |
| **p65** | Mouse | WB | WB 1:1000 | Cell Signaling Technology | 6956 |
| **p100/p52** | Rabbit | WB | WB 1:1000 | Cell Signaling Technology | 4882 |
| **Phospho-JNK** | Rabbit | WB | WB 1:500 | Cell Signaling Technology | 9255 |
| **Phospho-ERK** | Rabbit | WB | WB 1:1000 | Cell Signaling Technology | 9101 |
| **Phospho-p38** | Rabbit | WB | WB 1:500 | Cell Signaling Technology | 9216 |
| **p21** | Rabbit | WB | WB 1:2000 | Cell Signaling Technology | 2947 |
| **Vinculin** | Mouse | WB | WB 1:5000 | Sigma-Aldrich | V9131 |

**Table S9.**
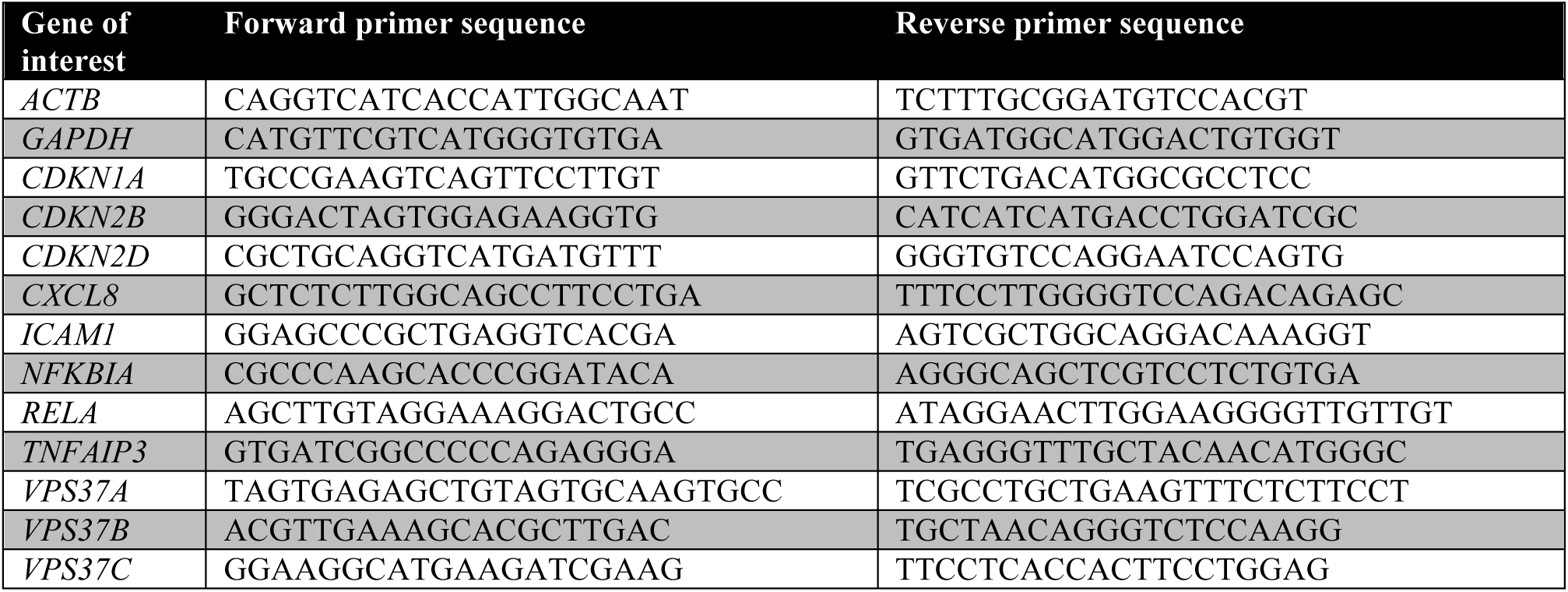
List of primers used in the present study.

## References

Aibar, S., Gonzalez-Blas, C. B., Moerman, T., Huynh-Thu, V. A., Imrichova, H., Hulselmans, G., Rambow, F., Marine, J. C., Geurts, P., Aerts, J. et al. (2017). SCENIC: single-cell regulatory network inference and clustering. Nat Methods 14, 1083–1086.

Alfred, V. and Vaccari, T. (2016). When membranes need an ESCRT: endosomal sorting and membrane remodelling in health and disease. Swiss Med Wkly 146, w14347.

Anders, S., Pyl, P. T. and Huber, W. (2015). HTSeq--a Python framework to work with high-throughput sequencing data. Bioinformatics 31, 166–9.

Bache, K. G., Slagsvold, T., Cabezas, A., Rosendal, K. R., Raiborg, C. and Stenmark, H. (2004). The growth-regulatory protein HCRP1/hVps37A is a subunit of mammalian ESCRT-I and mediates receptor down-regulation. Mol Biol Cell 15, 4337–46.

Banach-Orlowska, M., Jastrzebski, K., Cendrowski, J., Maksymowicz, M., Wojciechowska, K., Korostynski, M., Moreau, D., Gruenberg, J. and Miaczynska, M. (2018). The topology of the lymphotoxin beta receptor that accumulates upon endolysosomal dysfunction dictates the NF-kappaB signaling outcome. J Cell Sci 131.

Barbieri, E., Di Fiore, P. P. and Sigismund, S. (2016). Endocytic control of signaling at the plasma membrane. Curr Opin Cell Biol 39, 21–7.

Basile, J. R., Eichten, A., Zacny, V. and Munger, K. (2003). NF-kappaB-mediated induction of p21(Cip1/Waf1) by tumor necrosis factor alpha induces growth arrest and cytoprotection in normal human keratinocytes. Mol Cancer Res 1, 262–70.

Behan, F. M., Iorio, F., Picco, G., Goncalves, E., Beaver, C. M., Migliardi, G., Santos, R., Rao, Y., Sassi, F., Pinnelli, M. et al. (2019). Prioritization of cancer therapeutic targets using CRISPR-Cas9 screens. Nature 568, 511–516.

Bishop, N., Horman, A. and Woodman, P. (2002). Mammalian class E vps proteins recognize ubiquitin and act in the removal of endosomal protein-ubiquitin conjugates. J Cell Biol 157, 91–101.

Bonelli, P., Tuccillo, F. M., Borrelli, A., Schiattarella, A. and Buonaguro, F. M. (2014). CDK/CCN and CDKI alterations for cancer prognosis and therapeutic predictivity. Biomed Res Int 2014, 361020.

Brankatschk, B., Wichert, S. P., Johnson, S. D., Schaad, O., Rossner, M. J. and Gruenberg, J. (2012). Regulation of the EGF transcriptional response by endocytic sorting. Sci Signal 5, ra21.

Buser, D. P., Ritz, M. F., Moes, S., Tostado, C., Frank, S., Spiess, M., Mariani, L., Jeno, P., Boulay, J. L. and Hutter, G. (2019). Quantitative proteomics reveals reduction of endocytic machinery components in gliomas. EBioMedicine 46, 32–41.

Cai, Y., Crowther, J., Pastor, T., Abbasi Asbagh, L., Baietti, M. F., De Troyer, M., Vazquez, I., Talebi, A., Renzi, F., Dehairs, J. et al. (2016). Loss of Chromosome 8p Governs Tumor Progression and Drug Response by Altering Lipid Metabolism. Cancer Cell 29, 751–766.

Chen, F., Deng, J., Liu, X., Li, W. and Zheng, J. (2015). HCRP-1 regulates cell migration and invasion via EGFR-ERK mediated up-regulation of MMP-2 with prognostic significance in human renal cell carcinoma. Sci Rep 5, 13470.

Chen, F., Wu, J., Teng, J., Li, W., Zheng, J. and Bai, J. (2020). HCRP-1 regulates cell migration, invasion and angiogenesis via Src/ FAK signaling in human prostate cancer. Int J Biol Sci 16, 342–352.

Chen, F., Zhang, L., Wu, J., Huo, F., Ren, X., Zheng, J. and Pei, D. (2018). HCRP-1 regulates EGFR-AKT-BIM-mediated anoikis resistance and serves as a prognostic marker in human colon cancer. Cell Death Dis 9, 1176.

Colaprico, A., Silva, T. C., Olsen, C., Garofano, L., Cava, C., Garolini, D., Sabedot, T. S., Malta, T. M., Pagnotta, S. M., Castiglioni, I. et al. (2016). TCGAbiolinks: an R/Bioconductor package for integrative analysis of TCGA data. Nucleic Acids Res 44, e71.

Di Fiore, P. P. and von Zastrow, M. (2014). Endocytosis, signaling, and beyond. Cold Spring Harb Perspect Biol 6.

Dou, Y., Cha, D. J., Franklin, J. L., Higginbotham, J. N., Jeppesen, D. K., Weaver, A. M., Prasad, N., Levy, S., Coffey, R. J., Patton, J. G. et al. (2016). Circular RNAs are down-regulated in KRAS mutant colon cancer cells and can be transferred to exosomes. Sci Rep 6, 37982.

Du, Y., Wang, P., Sun, H., Yang, J., Lang, X., Wang, Z., Zang, S., Chen, L., Ma, J. and Sun, D. (2016). HCRP1 is downregulated in non-small cell lung cancer and regulates proliferation, invasion, and drug resistance. Tumour Biol.

El-Deiry, W. S. (2003). The role of p53 in chemosensitivity and radiosensitivity. Oncogene 22, 7486–95.

Floyd, S. and De Camilli, P. (1998). Endocytosis proteins and cancer: a potential link? Trends Cell Biol 8, 299–301.

Fu, F., Wan, X., Wang, D., Kong, Z., Zhang, Y., Huang, W., Wang, C., Wu, H. and Li, Y. (2018). MicroRNA-19a acts as a prognostic marker and promotes prostate cancer progression via inhibiting VPS37A expression. Oncotarget 9, 1931–1943.

Gargalionis, A. N., Karamouzis, M. V., Adamopoulos, C. and Papavassiliou, A. G. (2015). Protein trafficking in colorectal carcinogenesis-targeting and bypassing resistance to currently applied treatments. Carcinogenesis 36, 607–15.

Gingras, M. C., Kazan, J. M. and Pause, A. (2017). Role of ESCRT component HD-PTP/PTPN23 in cancer. Biochem Soc Trans 45, 845–854.

Gu, Z., Eils, R. and Schlesner, M. (2016). Complex heatmaps reveal patterns and correlations in multidimensional genomic data. Bioinformatics 32, 2847–9.

Guzman, C., Bagga, M., Kaur, A., Westermarck, J. and Abankwa, D. (2014). ColonyArea: an ImageJ plugin to automatically quantify colony formation in clonogenic assays. PLoS One 9, e92444.

Hayden, M. S. and Ghosh, S. (2008). Shared principles in NF-kappaB signaling. Cell 132, 344–62.

Hinata, K., Gervin, A. M., Jennifer Zhang, Y. and Khavari, P. A. (2003). Divergent gene regulation and growth effects by NF-kappa B in epithelial and mesenchymal cells of human skin. Oncogene 22, 1955–64.

Hoesel, B. and Schmid, J. A. (2013). The complexity of NF-kappaB signaling in inflammation and cancer. Mol Cancer 12, 86.

Hurley, J. H. (2015). ESCRTs are everywhere. EMBO J 34, 2398–407.

Keestra-Gounder, A. M., Byndloss, M. X., Seyffert, N., Young, B. M., Chavez-Arroyo, A., Tsai, A. Y., Cevallos, S. A., Winter, M. G., Pham, O. H., Tiffany, C. R. et al. (2016). NOD1 and NOD2 signalling links ER stress with inflammation. Nature 532, 394–7.

Krempler, A., Henry, M. D., Triplett, A. A. and Wagner, K. U. (2002). Targeted deletion of the Tsg101 gene results in cell cycle arrest at G1/S and p53-independent cell death. J Biol Chem 277, 43216–23.

Kwong, K. Y., Bloom, G. C., Yang, I., Boulware, D., Coppola, D., Haseman, J., Chen, E., McGrath, A., Makusky, A. J., Taylor, J. et al. (2005). Synchronous global assessment of gene and protein expression in colorectal cancer progression. Genomics 86, 142–58.

Lai, M. W., Huang, S. F., Lin, S. M., Chen, T. C., Lin, C. Y., Yeh, C. N., Yeh, T. S., Chen, M. F. and Yeh, C. T. (2009). Expression of the HCRP1 mRNA in HCC as an independent predictor of disease-free survival after surgical resection. Hepatol Res 39, 164–76.

Ledoux, A. C. and Perkins, N. D. (2014). NF-kappaB and the cell cycle. Biochem Soc Trans 42, 76–81.

Li, L. and Cohen, S. N. (1996). Tsg101: a novel tumor susceptibility gene isolated by controlled homozygous functional knockout of allelic loci in mammalian cells. Cell 85, 319–29.

Lin, Y. S., Chen, Y. J., Cohen, S. N. and Cheng, T. H. (2013). Identification of TSG101 functional domains and p21 loci required for TSG101-mediated p21 gene regulation. PLoS One 8, e79674.

Love, M. I., Huber, W. and Anders, S. (2014). Moderated estimation of fold change and dispersion for RNA-seq data with DESeq2. Genome Biol 15, 550.

Maminska, A., Bartosik, A., Banach-Orlowska, M., Pilecka, I., Jastrzebski, K., Zdzalik-Bielecka, D., Castanon, I., Poulain, M., Neyen, C., Wolinska-Niziol, L. et al. (2016). ESCRT proteins restrict constitutive NF-kappaB signaling by trafficking cytokine receptors. Sci Signal 9, ra8.

Manteghi, S., Gingras, M. C., Kharitidi, D., Galarneau, L., Marques, M., Yan, M., Cencic, R., Robert, F., Paquet, M., Witcher, M. et al. (2016). Haploinsufficiency of the ESCRT Component HD-PTP Predisposes to Cancer. Cell Rep 15, 1893–900.

Mattissek, C. and Teis, D. (2014). The role of the endosomal sorting complexes required for transport (ESCRT) in tumorigenesis. Mol Membr Biol 31, 111–9.

Meijer, G. A., Hermsen, M. A., Baak, J. P., van Diest, P. J., Meuwissen, S. G., Belien, J. A., Hoovers, J. M., Joenje, H., Snijders, P. J. and Walboomers, J. M. (1998). Progression from colorectal adenoma to carcinoma is associated with non-random chromosomal gains as detected by comparative genomic hybridisation. J Clin Pathol 51, 901–9.

Mellman, I. and Yarden, Y. (2013). Endocytosis and cancer. Cold Spring Harb Perspect Biol 5, a016949.

Mikula, M., Rubel, T., Karczmarski, J., Goryca, K., Dadlez, M. and Ostrowski, J. (2011). Integrating proteomic and transcriptomic high-throughput surveys for search of new biomarkers of colon tumors. Funct Integr Genomics 11, 215–24.

Miller, D. S. J., Bloxham, R. D., Jiang, M., Gori, I., Saunders, R. E., Das, D., Chakravarty, P., Howell, M. and Hill, C. S. (2018). The Dynamics of TGF-beta Signaling Are Dictated by Receptor Trafficking via the ESCRT Machinery. Cell Rep 25, 1841–1855 e5.

Moberg, K. H., Schelble, S., Burdick, S. K. and Hariharan, I. K. (2005). Mutations in erupted, the Drosophila ortholog of mammalian tumor susceptibility gene 101, elicit non-cell-autonomous overgrowth. Dev Cell 9, 699–710.

Morita, E., Sandrin, V., Chung, H. Y., Morham, S. G., Gygi, S. P., Rodesch, C. K. and Sundquist, W. I. (2007). Human ESCRT and ALIX proteins interact with proteins of the midbody and function in cytokinesis. EMBO J 26, 4215–27.

Mosesson, Y., Mills, G. B. and Yarden, Y. (2008). Derailed endocytosis: an emerging feature of cancer. Nat Rev Cancer 8, 835–50.

Nicolae, C. M., O’Connor, M. J., Constantin, D. and Moldovan, G. L. (2018). NFkappaB regulates p21 expression and controls DNA damage-induced leukemic differentiation. Oncogene 37, 3647–3656.

Olmos, Y. and Carlton, J. G. (2016). The ESCRT machinery: new roles at new holes. Curr Opin Cell Biol 38, 1–11.

Oughtred, R., Stark, C., Breitkreutz, B. J., Rust, J., Boucher, L., Chang, C., Kolas, N., O’Donnell, L., Leung, G., McAdam, R. et al. (2019). The BioGRID interaction database: 2019 update. Nucleic Acids Res 47, D529–D541.

Perisanidis, C., Savarese-Brenner, B., Wurger, T., Wrba, F., Huynh, A., Schopper, C., Kornek, G., Selzer, E., Ewers, R., Psyrri, A. et al. (2013). HCRP1 expression status is a significant prognostic marker in oral and oropharyngeal cancer. Oral Dis 19, 206–11.

Porther, N. and Barbieri, M. A. (2015). The role of endocytic Rab GTPases in regulation of growth factor signaling and the migration and invasion of tumor cells. Small GTPases 6, 135–44.

Rackov, G., Hernandez-Jimenez, E., Shokri, R., Carmona-Rodriguez, L., Manes, S., Alvarez-Mon, M., Lopez-Collazo, E., Martinez, A. C. and Balomenos, D. (2016). p21 mediates macrophage reprogramming through regulation of p50-p50 NF-kappaB and IFN-beta. J Clin Invest 126, 3089–103.

Sadler, J. B. A., Wenzel, D. M., Williams, L. K., Guindo-Martinez, M., Alam, S. L., Mercader, J. M., Torrents, D., Ullman, K. S., Sundquist, W. I. and Martin-Serrano, J. (2018). A cancer-associated polymorphism in ESCRT-III disrupts the abscission checkpoint and promotes genome instability. Proc Natl Acad Sci U S A 115, E8900–E8908.

Schmid, S. L. (2017). Reciprocal regulation of signaling and endocytosis: Implications for the evolving cancer cell. J Cell Biol 216, 2623–2632.

Schneider, C. A., Rasband, W. S. and Eliceiri, K. W. (2012). NIH Image to ImageJ: 25 years of image analysis. Nat Methods 9, 671–5.

Siegel, R. L., Miller, K. D. and Jemal, A. (2018). Cancer statistics, 2018. CA Cancer J Clin 68, 7–30.

Skrzypczak, M., Goryca, K., Rubel, T., Paziewska, A., Mikula, M., Jarosz, D., Pachlewski, J., Oledzki, J. and Ostrowski, J. (2010). Modeling oncogenic signaling in colon tumors by multidirectional analyses of microarray data directed for maximization of analytical reliability. PLoS One 5.

Stefani, F., Zhang, L., Taylor, S., Donovan, J., Rollinson, S., Doyotte, A., Brownhill, K., Bennion, J., Pickering-Brown, S. and Woodman, P. (2011). UBAP1 is a component of an endosome-specific ESCRT-I complex that is essential for MVB sorting. Curr Biol 21, 1245–50.

Stuchell, M. D., Garrus, J. E., Muller, B., Stray, K. M., Ghaffarian, S., McKinnon, R., Krausslich, H. G., Morham, S. G. and Sundquist, W. I. (2004). The human endosomal sorting complex required for transport (ESCRT-I) and its role in HIV-1 budding. J Biol Chem 279, 36059–71.

Sun, L., Lu, J., Ding, S., Bi, D., Ding, K., Niu, Z. and Liu, P. (2017). HCRP1 regulates proliferation, invasion, and drug resistance via EGFR signaling in prostate cancer. Biomed Pharmacother 91, 202–207.

Szymanska, E., Budick-Harmelin, N. and Miaczynska, M. (2018). Endosomal “sort” of signaling control: The role of ESCRT machinery in regulation of receptor-mediated signaling pathways. Semin Cell Dev Biol 74, 11–20.

Szymanska, E., Nowak, P., Kolmus, K., Cybulska, M., Goryca, K., Derezinska-Wolek, E., Szumera-Cieckiewicz, A., Brewinska-Olchowik, M., Grochowska, A., Piwocka, K. et al. (2020). Synthetic lethality between VPS4A and VPS4B triggers an inflammatory response in colorectal cancer. EMBO Mol Med 12, e10812.

Tanigawa, K., Maekawa, M., Kiyoi, T., Nakayama, J., Kitazawa, R., Kitazawa, S., Semba, K., Taguchi, T., Akita, S., Yoshida, M. et al. (2019). SNX9 determines the surface levels of integrin beta1 in vascular endothelial cells: Implication in poor prognosis of human colorectal cancers overexpressing SNX9. J Cell Physiol 234, 17280–17294.

Taniguchi, K. and Karin, M. (2018). NF-kappaB, inflammation, immunity and cancer: coming of age. Nat Rev Immunol 18, 309–324.

Tashima, T. (2018). Effective cancer therapy based on selective drug delivery into cells across their membrane using receptor-mediated endocytosis. Bioorg Med Chem Lett 28, 3015–3024.

Trakala, M., Arias, C. F., Garcia, M. I., Moreno-Ortiz, M. C., Tsilingiri, K., Fernandez, P. J., Mellado, M., Diaz-Meco, M. T., Moscat, J., Serrano, M. et al. (2009). Regulation of macrophage activation and septic shock susceptibility via p21(WAF1/CIP1). Eur J Immunol 39, 810–9.

Vasaikar, S., Huang, C., Wang, X., Petyuk, V. A., Savage, S. R., Wen, B., Dou, Y., Zhang, Y., Shi, Z., Arshad, O. A. et al. (2019). Proteogenomic Analysis of Human Colon Cancer Reveals New Therapeutic Opportunities. Cell 177, 1035–1049 e19.

Vietri, M., Radulovic, M. and Stenmark, H. (2020). The many functions of ESCRTs. Nat Rev Mol Cell Biol 21, 25–42.

Weinstein, J. N., Collisson, E. A., Mills, G. B., Shaw, K. R., Ozenberger, B. A., Ellrott, K., Shmulevich, I., Sander, C. and Stuart, J. M. (2013). The Cancer Genome Atlas Pan-Cancer analysis project. Nat Genet 45, 1113–20.

West, A. P., Khoury-Hanold, W., Staron, M., Tal, M. C., Pineda, C. M., Lang, S. M., Bestwick, M., Duguay, B. A., Raimundo, N., MacDuff, D. A. et al. (2015). Mitochondrial DNA stress primes the antiviral innate immune response. Nature 520, 553–7.

Wittinger, M., Vanhara, P., El-Gazzar, A., Savarese-Brenner, B., Pils, D., Anees, M., Grunt, T. W., Sibilia, M., Holcmann, M., Horvat, R. et al. (2011). hVps37A Status affects prognosis and cetuximab sensitivity in ovarian cancer. Clin Cancer Res 17, 7816–27.

Wood, L. D., Parsons, D. W., Jones, S., Lin, J., Sjoblom, T., Leary, R. J., Shen, D., Boca, S. M., Barber, T., Ptak, J. et al. (2007). The genomic landscapes of human breast and colorectal cancers. Science 318, 1108–13.

Wu, Y., Yang, Y. and Xian, Y. S. (2019). HCRP1 inhibits cell proliferation and invasion and promotes chemosensitivity in esophageal squamous cell carcinoma. Chem Biol Interact 308, 357–363.

Wuerzberger-Davis, S. M., Chang, P. Y., Berchtold, C. and Miyamoto, S. (2005). Enhanced G2-M arrest by nuclear factor-{kappa}B-dependent p21waf1/cip1 induction. Mol Cancer Res 3, 345–53.

Wunderley, L., Brownhill, K., Stefani, F., Tabernero, L. and Woodman, P. (2014). The molecular basis for selective assembly of the UBAP1-containing endosome-specific ESCRT-I complex. J Cell Sci 127, 663–72.

Xu, C. Y., Li, Z. J. and Hu, W. Z. (2017a). Up-regulation of HCRP1 inhibits proliferation and invasion in glioma cells via suppressing the ERK and AKT signaling pathways. Biomed Pharmacother 95, 31–36.

Xu, H., Miao, Z. F., Wang, Z. N., Zhao, T. T., Xu, Y. Y., Song, Y. X., Huang, J. Y., Zhang, J. Y., Liu, X. Y., Wu, J. H. et al. (2017b). HCRP1 downregulation confers poor prognosis and induces chemoresistance through regulation of EGFR-AKT pathway in human gastric cancer. Virchows Arch 471, 743–751.

Xu, J., Yang, W., Wang, Q., Zhang, Q., Li, X., Lin, X., Liu, X. and Qin, Y. (2014). Decreased HCRP1 expression is associated with poor prognosis in breast cancer patients. Int J Clin Exp Pathol 7, 7915–22.

Xu, J., Zhang, X., Wang, H., Ge, S., Gao, T., Song, L., Wang, X., Li, H., Qin, Y. and Zhang, Z. (2017c). HCRP1 downregulation promotes hepatocellular carcinoma cell migration and invasion through the induction of EGFR activation and epithelial-mesenchymal transition. Biomed Pharmacother 88, 421–429.

Xu, Z., Liang, L., Wang, H., Li, T. and Zhao, M. (2003). HCRP1, a novel gene that is downregulated in hepatocellular carcinoma, encodes a growth-inhibitory protein. Biochem Biophys Res Commun 311, 1057–66.

Xue, W., Kitzing, T., Roessler, S., Zuber, J., Krasnitz, A., Schultz, N., Revill, K., Weissmueller, S., Rappaport, A. R., Simon, J. et al. (2012). A cluster of cooperating tumor-suppressor gene candidates in chromosomal deletions. Proc Natl Acad Sci U S A 109, 8212–7.

Yang, W., Wang, J. G., Wang, Q., Qin, Y., Lin, X., Zhou, D., Ren, K., Hou, C., Xu, J. and Liu, X. (2016). Decreased HCRP1 promotes breast cancer metastasis by enhancing EGFR phosphorylation. Biochem Biophys Res Commun 477, 222–8.

Yang, W., Wang, J. G., Xu, J., Zhou, D., Ren, K., Hou, C., Chen, L. and Liu, X. (2017). HCRP1 inhibits TGF-beta induced epithelial-mesenchymal transition in hepatocellular carcinoma. Int J Oncol.

Yoshida, T., Kobayashi, T., Itoda, M., Muto, T., Miyaguchi, K., Mogushi, K., Shoji, S., Shimokawa, K., Iida, S., Uetake, H. et al. (2010). Clinical omics analysis of colorectal cancer incorporating copy number aberrations and gene expression data. Cancer Inform 9, 147–61.

Yu, G. and He, Q. Y. (2016). ReactomePA: an R/Bioconductor package for reactome pathway analysis and visualization. Mol Biosyst 12, 477–9.

Yu, G., Wang, L. G., Han, Y. and He, Q. Y. (2012). clusterProfiler: an R package for comparing biological themes among gene clusters. OMICS 16, 284–7.

Zhang, Q., Lenardo, M. J. and Baltimore, D. (2017). 30 Years of NF-kappaB: A Blossoming of Relevance to Human Pathobiology. Cell 168, 37–57.

Zhu, X., Liu, J., Xu, X., Zhang, C. and Dai, D. (2015). Genome-wide analysis of histone modifications by ChIP-chip to identify silenced genes in gastric cancer. Oncol Rep 33, 2567–74.

